# Organizational principles governing assembly and activation of the meiosis-specific Red1-Hop1-Mek1 complex

**DOI:** 10.1101/2022.04.06.487319

**Authors:** Linda Chen, Vaishnavi N. Nivsarkar, Saskia K. Funk, John R. Weir, Gerben Vader

**Affiliations:** Structural Biochemistry of Meiosis Group, Friedrich Miescher Laboratory, Tübingen, Germany; Department of Mechanistic Cell Biology, Max Planck Institute of Molecular Physiology, Dortmund, Germany; IMPRS for Living Matter, Max Planck Institute of Molecular Physiology, Dortmund, Germany; Amsterdam UMC location Vrije Universiteit, Section Oncogenetics, Department of Human Genetics, Amsterdam, The Netherlands; Cancer Center Amsterdam, Cancer Biology and Immunology, Amsterdam, The Netherlands; Amsterdam Reproduction & Development, Amsterdam, The Netherlands; Princess Máxima Center for pediatric oncology and Oncode Institute, Utrecht, The Netherlands

## Abstract

In mitosis, sister chromatids are preferred repair templates for homologous recombination, whereas in meiosis interhomolog-based repair is promoted. How this switch, which is a defining event in sexual reproduction, is accomplished remains poorly understood. In budding yeast, a meiosis-specific complex consisting of Red1, Hop1 and Mek1 (RHMc) enforces meiotic interhomolog bias, potentially through inhibition of intersister-based repair. The current data points to a linear assembly governing RHMc formation: the HORMA protein Hop1 associates with Red1 via a closure-motif-HORMA domain interaction, and Mek1 kinase is recruited through phospho-mediated interactions with Hop1. Here, via expression in mitotic cells we autonomously establish the RHM complex. *In vivo* analysis complemented with *in vitro* biochemical reconstitution shows that Mek1 associates with Red1, in a manner that might resemble binding of other kinases with scaffolding activators. The NH_2_-terminus of Red1 contributes to Hop1 binding, suggesting cooperative binding between Red1 and the HORMA domain of Hop1, beyond closure motif-based interactions. Meiotic activation of Mek1 kinase is dictated by complex formation and upstream DNA break-dependent signaling. We find Mek1 can be activated under DNA damaging conditions in mitotically dividing cells, where activation depends on upstream Mec1 kinase function and RHMc integrity. We perform a structure-function analysis of RHMc formation and Mek1 activation. Finally, we show that activation of Mek1 in mitosis leads to *rad51Δ*-like DNA break sensitivity, providing evidence for the model that RHMc instates meiotic interhomolog-based repair by inhibiting ‘mitotic’ homologous recombination. Our analysis enables querying downstream effects of RHMc action on DNA repair. Because aberrant re-expression of homologs of Red1 and Hop1 leads to DNA repair defects in human cancer, our system can be used to study roles of these genes during tumorigenesis.

## Introduction

Eukaryotes rely on meiosis to produce gametes (*i.e.,* sperm, egg or spores) that enable sexual reproduction. The biochemical principles that drive meiosis are similar to those fueling the canonical mitotic cell cycle, with additional meiosis-specific processes driving unique events required for gamete production^1^. Meiosis can thus be seen as an adaptation of the mitotic cell cycle. A hallmark of meiosis is a bias during homologous recombination (HR)-based DNA repair to use repair templates present on homologous chromosomes over those on sister chromatids ^2^. Such ‘interhomolog’ (IH) bias promotes repair of programmed DNA double strand breaks (DSBs) into interhomolog crossovers – a prerequisite for chromosome assortment and gamete production. This type of repair represents a remarkable adaptation of canonical HR repair that occurs in mitotically dividing cells, where sister chromatids are the preferred repair template during HR ^3^. Although factors have been described that promote interhomolog-based repair (such as meiosis-specific versions of the RecA recombinase (Dmc1) and several meiosis-specific auxiliary proteins that promote distant homology searches), how cells ‘inhibit’ HR via the proximal – and normally preferred – identical sequences present on sister chromatids remains a key question.

We know most about this step in meiotic HR from work in the budding yeast *Saccharomyces cerevisiae*. In this organism, the intersister-to-interhomolog template switch is controlled by the Red1-Hop1-Mek1 complex (*i.e.* the RHM complex; RHMc) (reviewed in ^2^) (**Figure 1a and b**). Without RHMc function, and Mek1 activity, meiotic DSB are effectively repaired using sister chromatids ^4–8^, leading to failed homolog linkage, impaired chromosome segregation and defective gamete formation ^9–12^. *RED1*, *HOP1* and *MEK1* are specifically expressed in meiosis, and they function together to establish IH bias ^13-19^). Red1 is a filamentous protein that is loaded onto meiotic chromosomes early in the meiotic program ^13,20–23^. Loss of Red1 *i)* disrupts meiosis-specific axis-loop chromosome organization ^24^, *ii)* impacts (Spo11-dependent) programmed DSB formation, *iii)* causes loss of interhomolog repair bias, and *iv)* leads to spore viability defects ^4,11,25,26^. Hop1 contains an NH_2_-terminal HORMA domain, which can topologically embrace a short peptide motif (termed ‘closure motif’; CM ^13,14^) present in Red1 ^13,12,22,23^ (**Figure 1a and b**). Hop1 is also a component of the meiotic chromosome axis ^27^ and required for efficient DSB formation ^12,25^. In addition to a HORMA domain ^28,29^, Hop1 contains a chromatin-binding domain ^30–32^ and a CM (*i.e.* a HORMA domain-binding peptide) at its COOH-terminus, which can be captured by Hop1’s own HORMA domain ^7,14,22,29^, potentially forming Hop1-to-Hop1 beads-on-a-string assemblies ^14^ or intramolecular CM-HORMA associations ^14,22,33^. Red1 and Hop1 are associated with ‘chromosome axis’ sites defined by meiotic (*i.e.* Rec8-containing) cohesin ^16,17^. Rec8-cohesin drives recruitment of Red1 and Hop1, potentially via a direct association ^17^, although a cohesin-independent Red1/Hop1 recruitment also occurs ^30,31^. Red1 and Hop1 recruit DSB factors to chromosome axis sites, likely via a direct association with Hop1 ^15–18,34–36^. Mek1, the third component of the RHMc, is a serine-threonine protein kinase that, in addition to its kinase domain, harbors a ForkHead-Associated (FHA) domain – a phospho-peptide binding moiety ^37,38,39^. Mek1 is related to Rad53, a key DNA damage checkpoint kinase ^38–40^ and homolog of the conserved CHK2 checkpoint kinase ^38–42^.

**Figure 1.**
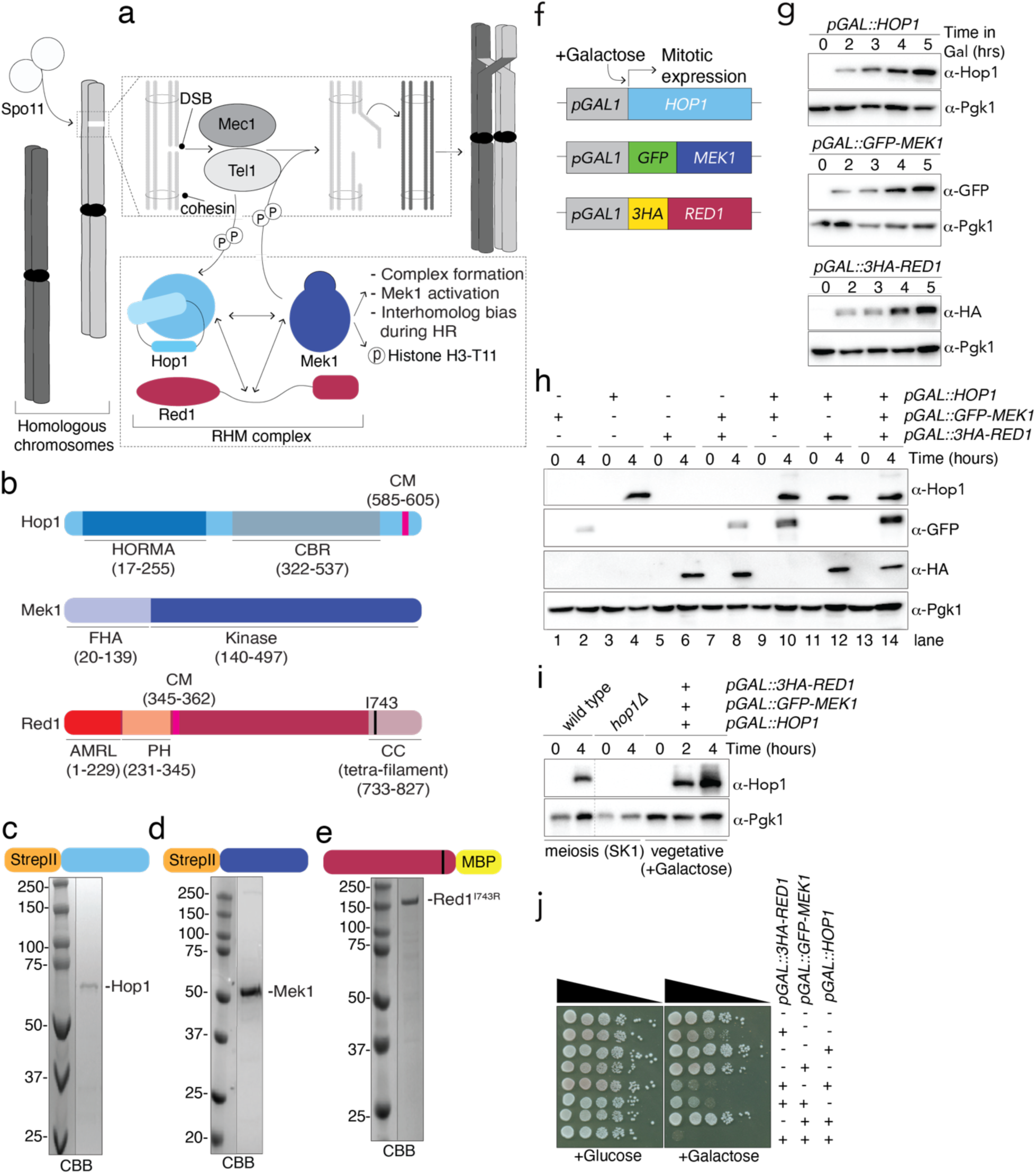
Expression of RHM complex subunits in mitosis and purified RHMc subunits. **a**. Schematic depicting the Red1-Hop1-Mek1 complex in meiosis, and the conceptual framework of the study. **b.** Schematic of domain organizations for Red1, Hop1 and Mek1. **c,d,e**. CBB stainings of purified proteins, as indicated. **f**. Schematic of galactose-inducible allele design for *RED1*, *HOP1* and *MEK1*. **g**. Expression analysis of Red1, Hop1, and Mek1 (strains used: yGV3726, yGV3243, and yGV2812). Galactose was added for the indicated time (hours). See also Supplementary Figure 1 for details on growth conditions. ⍺-HA was used to detect Red1, ⍺-GFP was used to detect Mek1, and ⍺-Hop1 was used to detect Hop1. Pgk1 was probed as loading control. **h**. Expression of Red1, Hop1, and Mek1 in mitotically-dividing cells in strains harboring *pGAL::3HA-RED1*, *pGAL::HOP1*, *pGAL::GFP-MEK1*, and combinations thereof. Strains used are yGV3726, yGV3243, yGV2812, yGV3235, yGV3255, yGV3219, and yGV4806. ⍺-HA was used to detect Red1, ⍺-GFP was used to detect Mek1, and ⍺-Hop1 was used to detect Hop1. Pgk1 was probed as loading control. **i**. Comparison of Hop1 expression in strains expressing Red1, Mek1 and Hop1 (yGV4806) in mitotic cells (induction time indicated) compared to Hop1 expression in meiotic (SK1) strains (wild type (yGV49) and *hop1Δ*(yGV4442)). In case of mitotic expression, galactose was added for the indicated time (hours). Samples taken at indicated times. ⍺-Hop1 was used to detect Hop1. Pgk1 was probed as loading control. **j**. Serial dilution (10-fold) spotting of yeast strains with indicated genotypes on Glucose- or Galactose-containing solid medium. Strains used are: yGV104, yGV3726, yGV3243, yGV2812, yGV3235, yGV3255, yGV3219, and yGV4806. **g**. Flow cytometry of wild type, *pGAL::3HA-RED1*, *pGAL::HOP1*, *pGAL::GFP-MEK1* and *RED1*, *HOP1* and *MEK1* strains, upon growth in glucose (+Glu) or galactose-containing (+Gal) conditions. Times (hours) indicated. DNA was stained with SYTOX Green. Strains used are: yGV104, yGV3726, yGV3243, yGV2812, and yGV4806.

Meiotic DSB formation triggers a signaling cascade known as the meiotic G2/prophase or pachytene checkpoint ^42,43^. Central to this checkpoint are sensor kinases, Tel1 and Mec1 – the budding yeast homologs of ATM and ATR, respectively – that also respond to mitotic DNA damage ^40,41^. Tel1 is activated at DSB sites, whereas Mec1 activation requires ssDNA tracts generated around DSB sites, through the action of defined nucleases. Mec1/Tel1 phosphorylate downstream substrates to coordinate cell cycle arrest with DNA repair. In mitosis, a central downstream kinase in the Mec1/Tel1 cascade is Rad53, whose activation relies on Mec1/Tel1-dependent phosphorylation of its adaptor protein, Rad9 ^44-45^. Normally, Rad53 is not activated in meiosis ^46^, and its role in the meiotic checkpoint is taken over by Mek1 ^46,47^. Hop1 plays a role that is conceptually similar to Rad9 for Mek1: it is a substrate of Mec1/Tel1, and its phosphorylation leads to chromosomal recruitment and activation of Mek1, likely via induced dimerization of Mek1 ^37,7,48–51^. Activated Mek1 phosphorylates several downstream substrates ^39,52–54^, as such influencing cell cycle arrest and DNA repair. By impinging on the meiotic transcription factor Ndt80, Mek1 halts meiotic cell cycle progression until DSB repair is complete ^39,52–54,55^. In addition, Mek1 activity promotes the establishment of IH repair bias ^6,56,57^. Mek1 substrates that impact DNA repair outcome have been identified, prominently among them Rad54 ^6,56,58^. Rad54 is an accessory factor to the DNA recombinase Rad51 ^58–62^, and phosphorylation of Rad54 inhibits the ability of Rad51 to promote homologous recombination ^6,56,58^.

A germane question is how an inhibitory effect on Rad51-dependent HR could lead to the promotion of interhomolog-biased repair. A model put forward by Subramanian and co-workers posits that this inhibitory effect – when Mek1 activity is restricted to the vicinity of DSB sites (and associated Mec1/Tel1 activity) – might uniquely prevent repair utilizing template on sister chromatids that reside in vicinity of the DSB (due to their close association brought about by sister chromatid cohesion) ^63^. Localized DSB repair inhibition could promote exploration towards distant repair template searches beyond the sphere of influence of localized Mek1 activity ^64^, eventually resulting in IH-based repair ‘activation’. Such a mechanism, that can distinguish repair templates based on spatial organization, would be conceptually reminiscent of the Aurora B kinase (Ipl1 in yeast)-dependent ‘spatial separation’ model that promotes kinetochore bi-orientation during mitotic chromosome segregation ^64,65^.

Connections between homologs through crossovers are a near-universal prerequisite for successful gamete formation, but little is known about the establishment of interhomolog HR bias outside of budding yeast ^2^. Nonetheless, homologs of Red1 and Hop1 are conserved ^2,66^, making it thus a distinct possibility that a complex that is functionally and molecularly analogous to the budding yeast RHM complex (possibly with CHK2 kinase as its enzymatic component ^67^, or partnered with a currently unknown functional Mek1 homolog) is important, also during human sexual reproduction.

To evaluate and critically interrogate the role of the RHMc in establishing IH-bias, it is pertinent to build a comprehensive molecular model of the assembly and activation principles of this complex, revealing catalytic mechanisms and regulatory control that can be linked to chromosomal events during meiotic G2/prophase. Due to the pleiotropic meiotic phenotypes associated with *RED1* and *HOP1* (see above), classical genetic approaches are inherently challenging to interpret. Alternative approaches to understand molecular systems are studies of assemblies in isolated, non-physiological environments, for example through *in vitro* biochemical reconstitutions (*e.g.,* for Spo11-mediated DSB formation ^68–71^), or via expression of factors outside their normal physiological setting (*e.g.*, expression of meiosis-specific factors in non-meiotic cells (*e.g., ^72–75^*). In this study, we develop both these experimental approaches to study molecular ‘rules of engagement’ governing RHMc formation and associated Mek1 activation.

## Results

To reveal the assembly principles of the RHMc complex we sought to establish conditions allowing isolation of individual components (and mutants thereof) from heterologous expression systems (See **Figure 1b** for schematic of individual RHM factors and key domain characteristics). The expression of recombinant Hop1 has been described by us and others ^15,22^. Utilizing an NH_2_-terminal twin Strep-II tag, we could affinity purify Hop1 followed by size exclusion chromatography (**Figure 1c)**. Inspection of the absorption at 260 and 280 nm indicated that the sample was free of nucleic acid contamination (**Supplementary Figure 1a**), and its elution pattern was consistent with a monomeric state. Mek1 was purified to homogeneity utilizing affinity capture on the NH_2_-terminal twin Strep-II tag (**Figure 1d and Supplementary Figure 1b and c**). Full-length, wild-type Red1 was not amenable to purification (data not shown), possibly due to the propensity of Red1 to form large homo-oligomeric filaments ^13^. To circumvent this, we mutated isoleucine 743 to arginine of Red1 (Red1^I743R^), the *Saccharomyces cerevisiae* equivalent to the I715R mutation in *Zygosaccharomyces rouxii* ^13^ (**Supplementary Figure 1d**), which was described to disrupt filament formation. Red1^I743R^-MBP could be purified to homogeneity (**Figure 1e**).

In parallel, we established a heterologous ‘*in vivo*’ system via expression of RHMc subunits in non-meiotic budding yeast cells. We placed the galactose-responsive *pGAL1* promoter in front of the coding regions of the genes encoding Red1, Hop1 and Mek1 – note that in case of *RED1* and *MEK1*, we added NH2-terminal affinity tags (3HA and GFP, respectively) to enable antibody-based detection (**Figure 1f**). When grown under conditions that lacked galactose, cells did not express *RED1*, *HOP1* or *MEK1*, and galactose addition led to rapid expression of RHMc subunits (**Figure 1g**). We generated strains expressing different combinations of the three *pGAL1*-regulated genes, and confirmed co-expression of *RED1*, *HOP1* and *MEK1* (**Figure 1h**). Protein levels of Hop1 upon mitotic *pGAL1*-driven expression were comparable with endogenous protein levels (*i.e*., in meiotic prophase cells, when RHMc subunits are expressed and functional) (**Figure 1i**). The levels of Hop1 in mitotically dividing cells increased further upon longer induction (*i.e*., after 4 hours). Thus – at least in the case of Hop1 – mitotic protein levels are comparable to those seen under physiological conditions in meiosis. Under certain allele combinations we noticed effects on protein stability – for example, Mek1 protein levels were consistently elevated when Hop1 and Red1 were also co-expressed (*e.g.*, compare ⍺-GFP (Mek1) signal in lane 2 and 14 of **Figure 1h**). This might reflect protein stabilization brought about by the presence of cognate binding partners, thus hinting at the possibility that Red1, Hop1 and Mek1 might associate under these conditions.

Mitotic cells expressing *RED1* experienced a growth defect (**Figure 1j**), which was aggravated by presence of *HOP1* and/or *MEK1* (**Figure 1j**). No defects were observed in cells expressing *HOP1* and/or *MEK1* in the absence of *RED1*. Flow cytometry revealed that *RED1* expression caused an increase in cells with a 2N DNA content, indicating a delay or arrest in G2/mitosis (**Supplementary Figure 2a**). These results are in line with reported effects of *RED1* overexpression on cell cycle progression ^76^. Cell cycle effects were exacerbated by co-expression of *HOP1* and *MEK1* (**Supplementary Figure 1c**), mirroring viability defects (**Figure 1j**). We speculate that effects on cell growth might be related to Red1 higher-order assemblies – specifically tetramers as well as filaments of tetramers ^13,12,20–23^.

We looked for interactions between Red1, Hop1 and Mek1, as occurs in meiosis. Indeed, immunoprecipitation of Red1 led to enrichment of Mek1 or Hop1 (**Figure 2a**). A similar association between RHM components was detected upon immunoprecipitation of GFP-Mek1 (**Supplementary Figure 2b**). Thus, the RHM complex can autonomously be established in mitotic cells. These data suggest that, in addition to its described interaction with Hop1 (via its closure motif (CM) encoded in Red1^340-362^ (**Figure 1b**); see also below ^12,13,20,23^, Red1 encodes a (Hop1-independent) Mek1-interaction region. This was surprising, since RHMc assembly is often portrayed as a linear series of events (*e.g*.,^6,7,12,13,20,23^), in which Red1 first associates with Hop1 (mediated by HORMA-CM association) ^12,22^. Subsequent association of Mek1 with Hop1, mediated by FHA-domain based association with phosphorylated Hop1 (through Mec1/Tel1-dependent phosphorylation of Hop1) would establish trimeric RHMc formation ^7,48,49,51^. We sought to understand this interaction in more detail. The NH_2_-terminus of Red1 encodes a predicted folded domain (amino acids 1-340). Analysis of the AlphaFold2 structure of Red1 (https://alphafold.ebi.ac.uk/entry/P14291) revealed similarity of this region to the structure of the human meiotic protein SYCP2, a Red1 ortholog ^77^ (**Figure 2b**). Red1 and SYCP2 both harbor an NH_2_-terminal domain encoding an Armadillo Repeat Like (ARML) domain followed by a second folded domain – originally coined as an Spt16M-like-domain (SLD) based on the structural similarity to the middle-region of Spt16 ^77^, but more generally described as a Pleckstrin Homology (PH) domain ^66,77,78^. We will here refer to this region as the PH domain ^66,77,78^ (**Figure 1b** and **2b**, and see below). The AMRL-PH module is upstream of the Red1 closure motif sequence (CM, amino acids 341-362; magenta in Red1 schematic in **Figure 1b, 2b**)) that mediates interaction with Hop1 via CM-HORMA domain binding ^12,22^. This region is followed by a large (likely unstructured) region (362-362) and a COOH-terminal coiled-coil region (Red1^340-362^) ^12,13,20^. This coiled-coil domain of Red1 harbors tetramerization as well as filament forming activities in other yeasts ^12,13,20^. Filamentous Red1 assemblies are thought to be crucial to establish the meiotic chromosome structure ^12,13,20^. Mutations that disrupt this filament-forming domain lead to defects in meiotic chromosome organizations and spore viability ^79^. Truncation of the extreme COOH-terminal region of Red1 (*i.e*., removal of the last 7 amino acids, 820-827) was shown to lead to a specific disruption of filament formation, while leaving tetramerization unaffected (^12,13,20^). We generated truncation alleles of Red1 (all driven by *pGAL-3HA)*, based on these structural features of Red1 (**Figure 2c**). We performed co-immunoprecipitation analysis to investigate the requirement of Red1 to interact with Hop1 and/or Mek1 (**Figure 2d**). As expected, the *in vivo* interaction between Hop1 and Red1 depended on the presence of the CM within Red1 ^12,22^ : a construct of a Red1 expressing the first 345 amino acids but lacking the CM failed to co-purify Hop1 (Red1^1-345^; **Figure 2c and d**). A Red1 fragment that included the CM of Red1 (346-827) was able to interact with Hop1 (**Figure 2e**), whereas a larger truncation which removed the CM of Red1 (346-827) failed to bind to Hop1. Full length Red1 (thus also encoding the structured ARM/PH module directly adjacent to the CM) bound more efficiently to Hop1 as compared to the version of Red1 that was truncated upstream of the CM (**Figure 2e**). This suggests that the domain organization of Red1, in which the CM is located immediately adjacent to a structured region consisting of ARML-PH, might influence Hop1-Red1 association. Thus, the region of Red1 upstream of the CM could contribute to Hop1 binding directly. Alternatively, the structured NH_2_-terminal domain might prevent ‘sliding off’-based dissociation of CM in Red1 from the HORMA domain of Hop1, reminiscent of what was recently described for the association between the HORMA domain of Mad2 and its topological binding partner Cdc20 ^80^.

**Figure 2.**
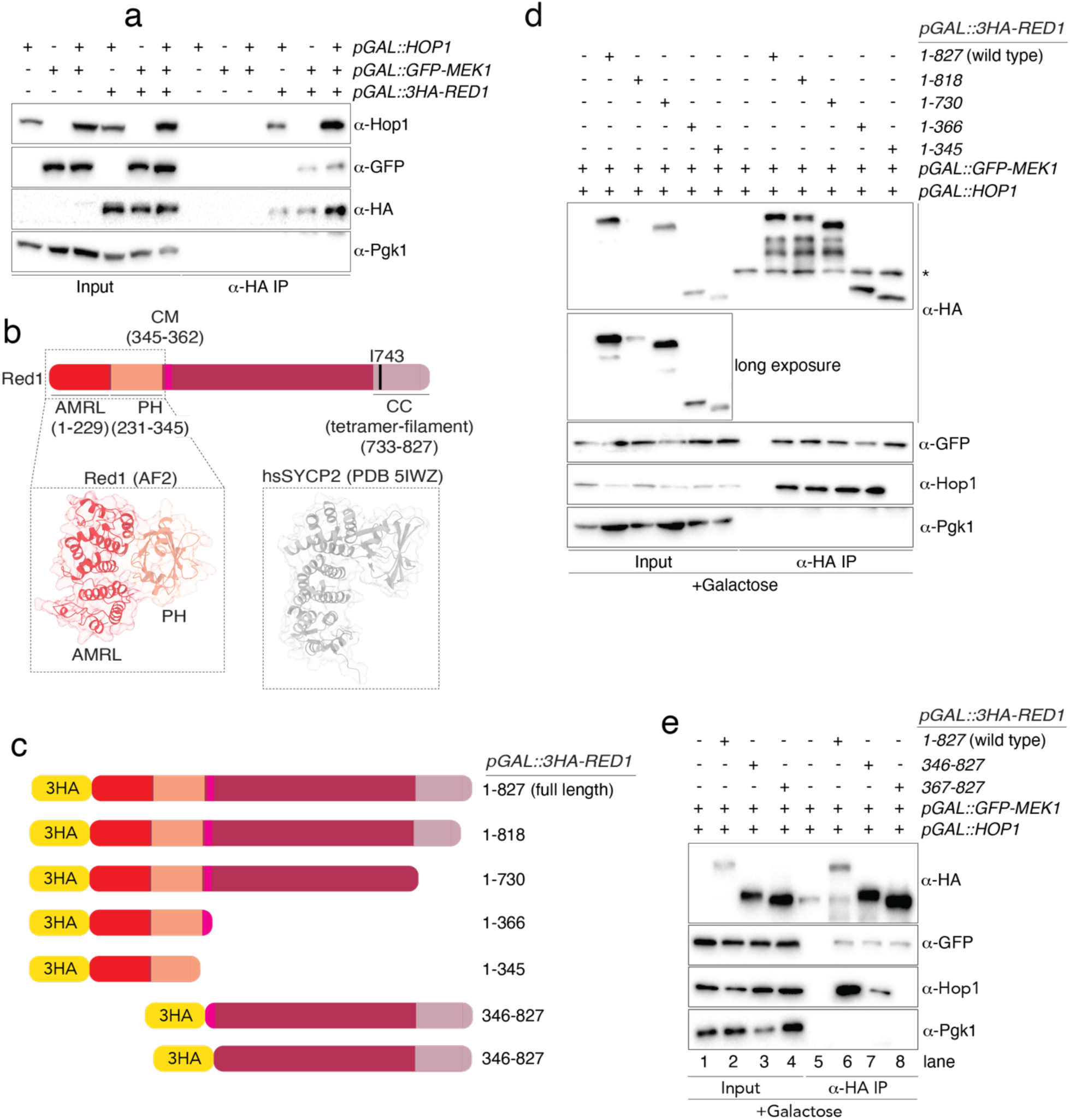
Red1-Hop1-Mek1 complex formation in mitosis. **a**. Co-immunoprecipitation (co-IP) between Red1, Hop1 and/or Mek1. Red1 was immunoprecipitated via ⍺-HA pulldown. ⍺-HA was used to detect Red1, ⍺-GFP was used to detect Mek1, and ⍺-Hop1 was used to detect Hop1. Pgk1 was probed as loading control. Samples were taken after 4 hours induction with galactose. The following strains were used: yGV3242, yGV2812, yGV3235, yGV3255, yGV3219, yGV4806. **b**. Schematic of Red1, with domain structure and AlphaFold-based model of Red1^ARML-PH^ compared to the structure of the NH_2_-terminus of human SYCP2 (PBD 5IWZ). **c**. Schematic depicting Red1 truncation mutants employed in this figure/study. **d**. Co-immunoprecipitation (co-IP) between Red1 (wild type and truncations), and Hop1 and/or Mek1. Red1 was immunoprecipitated via ⍺-HA pulldown. ⍺-HA was used to detect Red1, ⍺-GFP was used to detect Mek1, and ⍺-Hop1 was used to detect Hop1. Pgk1 was probed as loading control. * indicates IgG heavy-chain. Samples were taken after 4 hours induction with galactose. The following strains were used: yGV3219, yGV4806, yGV4393, yGV4395, yGV4397, yGV4400. **e**. Co-immunoprecipitation (co-IP) between Red1 (wild type and truncations), and Hop1 and/or Mek1. Red1 was immunoprecipitated via ⍺-HA pulldown. ⍺-HA was used to detect Red1, ⍺-GFP was used to detect Mek1, and ⍺-Hop1 was used to detect Hop1. Pgk1 was probed as loading control. Samples were taken after 4 hours induction with galactose. The following strains were used: yGV3219, yGV4806, yGV4207, yGV4402.

We next focused on the incorporation of Mek1 into the RHMc. A fragment of Red1 that contains its NH_2_-terminal domain (1-345) was able to pull down Mek1 (**Figure 2d**), whereas truncated Red1 fragments that lacked this domain but contained the COOH-terminal part of Red1 (*e.g.*, Red1^346-827^ or Red1^367-827^) were also proficient for Mek1 interaction (**Figure 2e**). Associations occurred independently of Hop1 presence (see pulldowns for Red1 fragments 1-345 and 367-827 in **Figure 2d and e**), suggesting that direct Red1-Mek1 associations can be established. Thus, at least under the mitotic expression conditions, Red1 contains multiple (independent) binding sites for Mek1, which do not require the presence of Hop1.

The assembly principles of the RHM complex were next studied using recombinant protein approaches, with the purified proteins described in **Figure 1c-e**. In addition to the production of full-length proteins, we produced numerous protein fragments and mutations which correspond to the putative domains of the RHMc subunits. Our findings using co-IP in mitotic yeast cells (**Figure 2, and Supplementary Figure 2**) hint at the existence of a direct, potentially composite mode of association between Red1, Mek1 and Hop1. We produced four constructs of Red1 containing amino acids 1-230 (corresponding to the ARML domain), 230-345 (for the PH domain) and 1-362 (the ARML-PH domains and the closure motif (CM)) of Red1 (**Figure 3a**). Due to the difficulty in purifying all of these recombinant proteins we utilized a co-expression approach in insect cells, where these different NH_2_-MBP-tagged constructs were co-expressed with 2xStrep-II tagged Mek1. Affinity purification via the 2xStrep-II tag, revealed that all Red1 constructs showed (differing) ability to associate with Mek1. These data indeed confirm that the NH_2_-terminal part of Red1 can associate with Mek1. The most prominent association was found for Red1^230-345^ (**Figure 3b**). We employed predictive tools to determine if we could derive a plausible model for the Red1-Mek1 interaction which would be in agreement with our *in vitro* experimental analysis. Attempts with full-length proteins were not fruitful, so we turned to shorter fragments of both Red1 and Mek1. The best quality prediction we could obtain suggested a direct interaction between the ARML domain of Red1 (1-230) and the kinase domain of Mek1 (**Figure 3c and d**), which in principle is in agreement with interactions detected in our *in vitro* and yeast analysis. Sequence conservation mapping onto the surface of the Red1^ARML^ domain reveals that the region predicted to form a binding interface with Mek1 kinase is indeed highly conserved (**Supplementary Figure 3a-b**). Strikingly however, in our prediction, the potential kinase binding site present in this region of Red1 is occupied by the Red1^PH^ domain (residues 227-345) (compare **Figure 2b** to **3f**)). This arrangement is not compatible with the position of the PH domain of Red1 in the AlphaFold2 model (due to a steric clash between PH domain and the kinase domain of Mek1), nor with the experimentally determined structure of SYCP2 (**Figure 2b**). Therefore, if the AlphaFold2 model of the Red1-Mek1 complex is correct, there must be movement of the Red1^PH^ domain relative to the Red1^ARML^ domain to accommodate Mek1 association (**Figure 3e**). We note that in our *in vitro* pull-down experiments we detected a robust interaction between the PH-domain of Red1 (230-345) and the full length Mek1(**Figure 3b**), which might point to a multistep association cascade in which Red1^PH^-Mek1 interactions cause a displacement of the PH domain, enabling the establishment of the Red1^ARML^-Mek1 kinase domain assembly as predicted by structural modeling. In either case, our data using *in vitro* reconstitutions and *in silico* modeling indeed points to the existence of a direct interaction between Red1 and Mek1, as suggested by our experiments in mitotic budding yeast (**Figure 2**). We compared the predicted structure of these domains of experimentally determined structures available in the PDB using DALI ^81^. The PH domain of Red1 showed high similarity (Z-score of 9.0 with a C⍺ RMSD of 2.3Å over 125 residues) to the Red1^PH^ domain of mouse REC114 ^82^, hinting at a common origin of these two meiotic recombination factors. The ARML domain of Red1 was found by a DALI search to be structurally highly similar to MO25b (PDB 3ZHP 81 with a C⍺ RMSD of 2.9 Å (over 152 residues)^82,83^, **Supplementary Figure 4a**). Significantly, MO25b is a kinase adapter protein that enables kinase activation ^84^. We used the structure of the STK24 kinase with MO25b^82,83^ to superimpose Mek1 onto the STK24 kinase (C⍺ RMSD of 2.6 Å (over 233 residues)), **Supplementary Figure 4b**), which revealed a similar position relative to MO25b/Red1-ARML. Again, these analyses point to a potential mode of interaction between Red1 and Mek1 that is structurally similar to that of MO25B and STK24, with the important distinction that accommodating this kind of binding necessitates significant spatial movements of the Red1^PH^ domain to allow binding to the Red1^ARML^ domain with Mek1 (**Figure 3e**)

**Figure 3.**
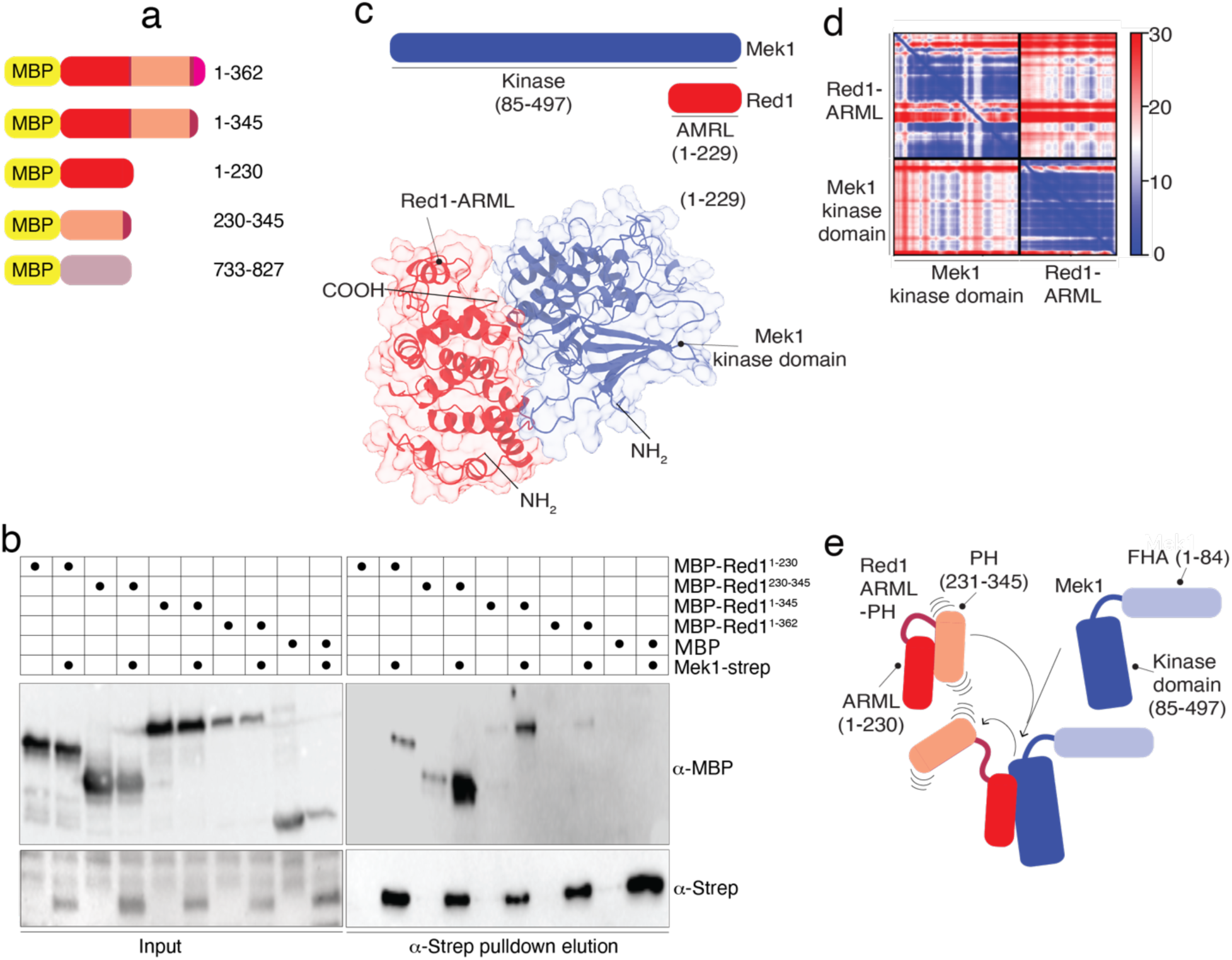
*In vitro* analysis of the Red1-Mek1 interaction. **a**. Schematic depicting Red1 truncation mutants employed in this figure/study. B. Pulldown experiments (⍺-Strep-based) of Mek1 together with indicated Red1 truncations, as shown in a. ⍺-Strep was used to detect Mek1, and ⍺-MBP was used to detect Red1 fragments. **c**. and **d**. AlphaFold-based modeling of Red1^ARML^-Mek1^kinase^ ^domain^ association, including Predicted Alignment Error (PAE) plots for this model. Red and blue coloring corresponds to high-confidence-to-low-confidence distributions. **e**. Schematic of a speculative model regarding dynamic association mode between Red1^ARML-PH^-Mek1.

We extended our studies to longer Red1 constructs. As described above, we had difficulty producing full-length recombinant Red1. We utilized both a point mutant (I743R), and a C-terminal truncation (ending at residue 819) analogous to those previously described to prevent filament formation of Red1, but retain tetramerization of the C-terminal coiled-coils ^13,20^ (**Figure 4a** and **Supplementary Figure 1d**). We co-expressed full-length Mek1 with MBP-tagged Red1, Red1^I743R^, Red1^1-819^ and Red1^1-819/I743R^. The expression levels of the Red1^1-819^ and full length Red1 were relatively low, so we excluded these from further analysis. Despite the expression levels of Red1^I743R^ and Red1^1-819/I743R^ being similar we only observed a robust interaction between Mek1 and Red1^1-819/I743R^ (**Figure 4b**, compare lane 4 and 8). Based on the work by Corbett and co-workers ^13^, the COOH-terminal truncation of Red1 is expected to behave the same as the I743R mutant – *i.e.*, it should remain tetrameric but not form filaments (**Figure 4a)**. Thus, our data might indicate that filament formation has a negative effect on the affinity of Mek1 for Red1. We focused on the Red1 coiled-coil containing regions in more detail. We purified different Red1 COOH-terminal coiled-coiled (CC) domains: Red1 733-827^WT^, 733-827^I743R^, 733-819^WT^ and 733-819^I743R^ using amylose affinity chromatography followed by size-exclusion. We analyzed Red1^733-827^ and Red1^733-827/I743R^ by SEC-MALS (**Figure 4c**). Consistent with previous observations, Red1^733-827^ formed large assemblies (estimated size ∼ 4 MDa; monomer size is 54,4 kDa). As expected ^13^, analysis of the MBP-Red1^733-827/I743R^ mutant by SEC-MALS showed the formation of species corresponding to tetramers (**Figure 4c**, estimated size ∼ 200-250 kDa; monomer size is 54,4 kDa)). In these SEC-MALS experiments we utilized protein samples at a concentration of 10 µM. We next utilized mass photometry to measure the mass of MBP-Red1 fragments at much lower concentrations of ∼100 nM (**Figure 4d**). Under these conditions the MBP-Red1^733-827/I743R^ formed species consistent with a dimer, rather than tetramers as was the case for Red1^733-819^ and Red1^733-819/I743R^. MBP-Red1^733-827^ formed species consistent with dimers and tetramers. Based on these observations we conclude that the Red1 coiled-coil domain forms concentration-dependent oligomers. We hypothesize that the difference in Red1 versus Red1^I743R^ in association with Mek1 (**Figure 4b**) might derive from the fact that Red1^733-819/I743R^ dissociates more readily into dimers. Thus, association of Red1 with Mek1 might be negatively influenced by tetramerization of Red1, at least under these *in vitro* concentrations and conditions (**Figure 4e**).

**Figure 4.**
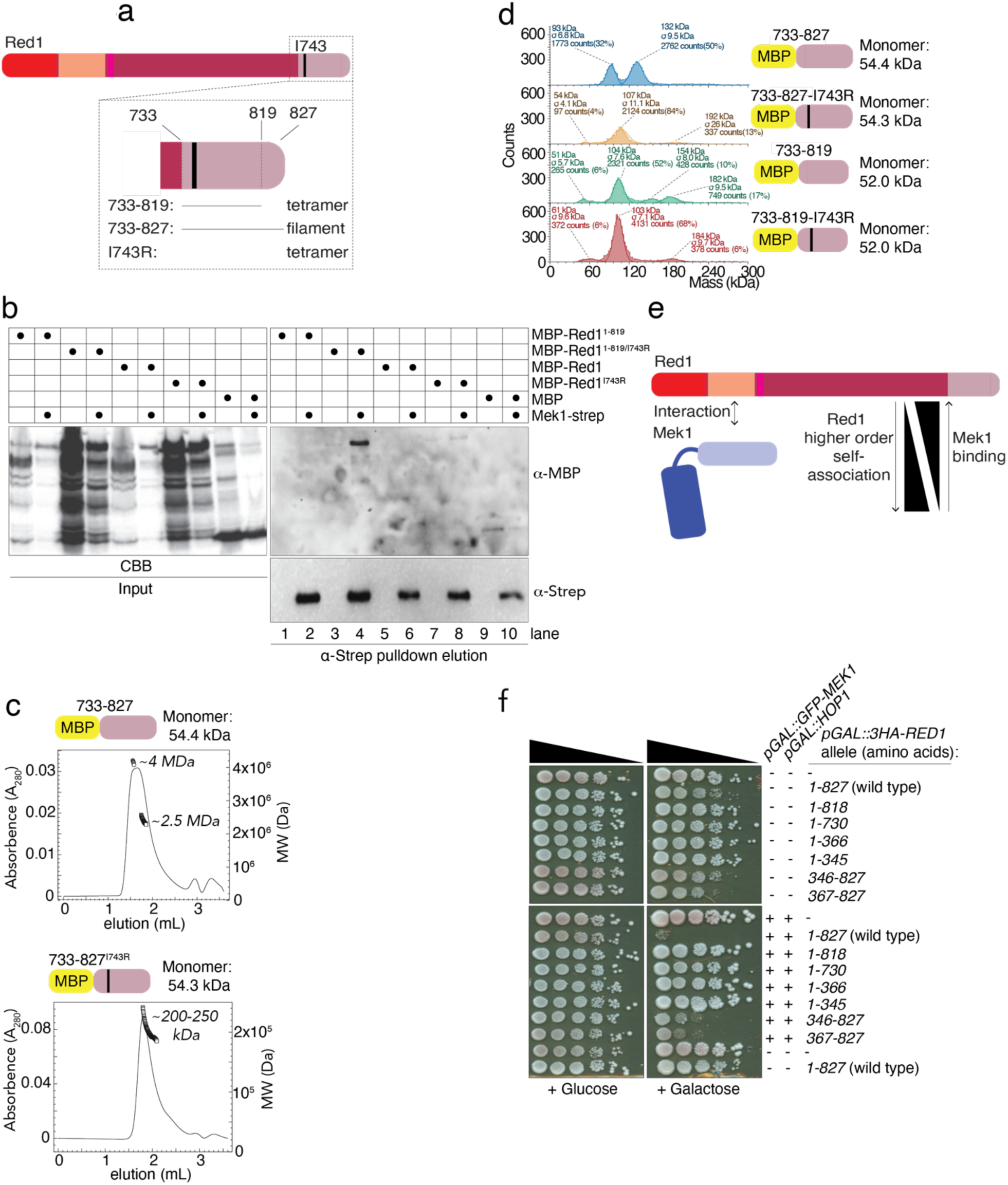
*In vitro* analysis of the Red1-Mek1 interaction and higher-order structural assemblies of Red1. **a**. Schematic depicting Red1 truncation mutants employed in this figure/study. **b**. Pulldown experiments (⍺-Strep-based) of Mek1 together with indicated Red1 truncations, as shown in a. ⍺-Strep was used to detect Mek1, and ⍺-MBP was used to detect Red1 fragments. CBB = Coomassie brilliant blue. **C**. SEC-MALS analysis of MBP-Red1 (733-827) and MBP-Red1 (733-819) at 10 µM. d. Mass photometry analysis of the Red1 coiled-coil protein truncations at 100 nM. Peaks were fitted by Gaussian curves in DiscoverMP. **e**. Schematic of speculative model regarding correlation between Mek1-binding propensity and higher-order self-association of COOH-terminus of Red1. **f**. Serial dilution (10-fold) spotting of yeast strains expressing different Red1 truncations with and without Hop1 and Mek1, on Glucose- or Galactose-containing solid medium. Strains used were: yGV104, yGV3726, yGV3219, yGV3798, yGV3799, yGV4190, yGV4191, yGV4193, yGV4194, yGV4207, yGV4393, yGV4395, yGV4397, yGV4400, yGV4402 and yGV4806.

We correlated this Red1 truncation-interaction analysis with our earlier findings that expression of full length Red1 (in isolation, or in combination with Hop1 and/or Mek1) led to growth defects (**Figure 1j**). We expressed different Red1 constructs, and queried effects on cell growth **(Figure 4f**). Expression of Red1 fragments that contained the extreme COOH-terminal amino acid stretch, known to lead to filament/tetramer formation, led to cell growth that were comparable to those seen upon the expression of full length Red1. Removing the filament-forming amino stretch (*i.e.*, in Red1^1-818^) abrogated these effects ^13,20^, hinting that the observed phenotypes on cell cycle progression and growth are linked to Red1 filament formation.

Hop1 interacts with Red1 in a manner dependent upon the closure motif of Red1 (residues 340-362) ^22^. We sought to gain further structural insights into this interaction using AlphaFold2-Multimer ^37,85^(**Figure 5a and b**). To our surprise the model suggested that there could be a direct interaction between the Red1 NH_2_-terminal domains and Hop1^HORMA^ independent of the closure motif (see also **Supplementary Figure 5a**).

**Figure 5.**
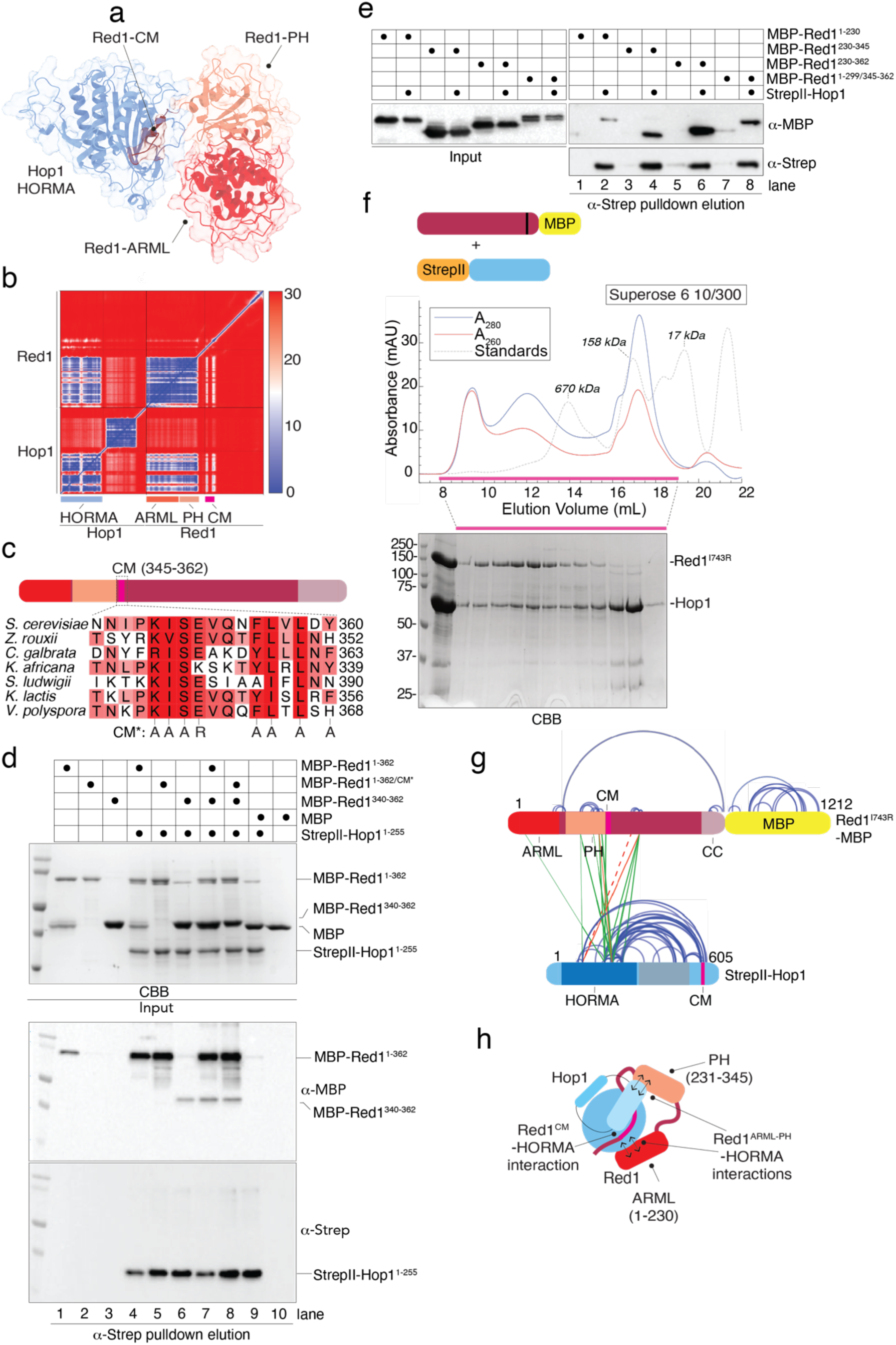
*In vitro* analysis of the Red1-Hop1 interaction. **a**. and **b**. AlphaFold-based modeling of Red1^ARML-PH^-Hop1^HORMA^ association, including Predicted Alignment Error (PAE) plots for this model. **c**. Sequence alignment of CM-region in Red1, including mutated residues in Red1^CM*^ mutant. **d.** and **e**. Pulldown experiments (⍺-Strep-based) of Hop1 together with indicated Red1 truncations, as shown in a. ⍺-Strep was used to detect Mek1, and ⍺-MBP was used to detect Red1 fragments. CBB = Coomassie brilliant blue. **f**. SEC run, including SDS-PAGE, of Red1^I743R^-MBP and StrepII-Hop1. CBB = Coomassie brilliant blue. **g.** XL-MS map of Red1^I743R^-MBP and StrepII-Hop1. Cross-links were filtered for a match score of >100 leading to an FDR of <1%. Intramolecular cross links are shown in blue. Intermolecular cross-links are colored according to whether they are consistent with the model (see main text).

We produced a mutant version of the closure motif, where eight conserved residues are mutated into either alanine or lysine from here on referred to as CM* (**Figure 5c**). We first tested whether this mutation was sufficiently penetrant by utilizing the wild-type or mutant closure motif fused to an N-terminal MBP domain in a pulldown experiment against Hop1. Note that for this experiment we used a version of Hop1 which carries a described mutation in its COOH-terminal CM (Hop1^K593A^) ^22,7^, to prevent self-closure of the HORMA domain. Indeed, only the wild-type CM sequence could capture Hop1, with no detectable binding to the MBP-CM* entity (**Supplementary Figure 5b**). Having confirmed our CM mutations, we next tested whether NH_2_-terminal Strep-tagged Hop1^HORMA^ could capture Red1^1-362/CM*^, as would be expected based on our in *silico* modeling. In a pulldown experiment we indeed observed interactions between Hop1 and Red1^1-362^ and Red1^1-362/CM*^ (**Figure 5d**, lanes 4 and 5), thus confirming our model in which a Hop1 to Red1 interaction can take place independently of the CM sequence. To test this further we asked what would happen if we separately added the Red1-CM (as MBP Red1^340-362^) to the pulldown. If the binding between Hop1-Red1 can indeed be established independently of a CM-mediated interaction, adding an ‘external’ CM should not interfere with binding. Indeed, binding of the external CM to Hop1 was fully compatible with Hop1 interaction with binding to Red1^1-362^ (**Figure 5d**, compare lanes 4 and 5 to 7 and 8), again lending support to the existence of a CM-independent binding interface between Hop1 and Red1. Based on the model, we would expect to see a significant loss of binding affinity for Hop1 when the closure motif of Red1 was mutated, however in pulldowns both Red1 1-362 and Red1 1-362 CM* showed similar apparent affinity for Hop1. This, while CM binding does impact the binding affinity of the Red1-Hop1 association, our data suggests that under these binding conditions the NH_2_-terminal domains of Red1 provide a sufficient affinity for Hop1.

Next, we further dissected the NH_2_-terminus of Red1 and its role in Hop1 binding. We made use of the same Red1 N-terminal fragments as described above. In a co-expression experiment from insect cells, we observed some interaction between Red1^1-230^ and Red1^230-345^ and Hop1, with considerably more Red1^230-345^ being pulled down on Hop1 (**Figure 5e**, compare lane 2 and 4). The inclusion of the closure motif alone was sufficient to robustly bind to Hop1 regardless of the rest of the Red1 sequence that was included (**Figure 5e**, lane 6). Despite the apparent congruence between the AlphaFold2 model of Hop1 and Red1 and our pulldown experiments, we considered further interactions between Hop1 and Red1. We produced a complex of full-length Hop1 with MBP-Red^1-827^ with the I743R mutation. The initial affinity purification of the complex showed apparently reasonable purity (**Figure 5f**). We evaluated the size and stoichiometry of the complex using mass photometry (**Supplementary Figure 5c**). The largest species we observed was determined at ∼209 kDa, which corresponds well to a 1:1 complex of Hop1 and Red1^I743R^ (theoretical molecular mass of 212 kDa). Unlike in the experiments with the N-terminally MBP-tagged Red1 coiled-coil constructs (residues 733-827 with I743R mutation) we did not observe species that corresponded to a Red1 dimer. We hypothesize that this is either due to the even lower relative concentration of Red1 or that the presence of the COOH-terminal MBP tag in this experiment interferes with oligomerization of Red1. We observe masses that correspond to both free monomeric Hop1 and free monomeric Red1^I743R^ (∼81 kDa and ∼140 kDa respectively; theoretical mass 71 kDa and 138 kDa, respectively). As such, we conclude that a 1:1 Hop1-Red1^I743R^ complex can form *in vitro*, but that this complex has partly dissociated at the low concentration (30 nM) of complex employed for mass photometry. We note that we also have an excess of Hop1 in our preparations (**Supplementary Figure 5d**). We subjected the Hop1-Red1^I743R^ complex to cross-linking with the bifunctional cross-linker DSBU, followed by proteolytic digestion and mass spectrometry, as described previously ^86^ (**Figure 5g**). Analysis of obtained cross-links revealed that Hop1 appears to be more extensively cross-linked than Red1^I743R^-MBP, likely reflecting excess free Hop1 in our purification. In Red1 ^I743R^-MBP, we observed a single long-distance cross-link between the COOH-terminal coiled-coil domain and the Red1^ARML/PH^ domains, which could conceivably either be an inter- or intramolecular crosslink.

A central functionality of the RHM complex in mediating DNA repair template decisions (and checkpoint function) lies in the kinase activation of Mek1, which leads to downstream phosphorylation events ^52–54,87^. The establishment of the RHM complex outside of its ‘natural’ environment prompted us to investigate whether, in mitotically dividing cells, this situation could be associated with activation of Mek1. To evaluate Mek1 kinase activity, we monitored the phosphorylation of Threonine 11 on Histone H3 (phospho-Histone H3-T11), a well-characterized substrate of Mek1 during meiotic prophase ^87,88^. We note that we occasionally observed an apparent background level of phospho-Histone H3-T11 in our mitotic culture conditions (*e.g*., see * in **Figure 6a**), possibly reflecting modification of Histone H3-T11 through a (Mek1-independent) pathway that is activated under certain nutritional conditions ^89,90^. We initially tested whether expression of Mek1 alone would lead to kinase activation. We found that mere expression of *GFP-MEK1* was not associated with an increase in Histone H3-T11 phosphorylation (**Figure 6a**). In meiosis, Mek1 activity is coupled to upstream Mec1/Tel1-dependent phospho-signaling triggered by Spo11-dependent DSB formation ^48,49^, a signaling module that is obviously lacking in our mitotic system. We next thus queried whether inducing DNA damage in combination with Mek1 expression led to its activation. We treated cells with methyl methanesulfonate (MMS), a DNA alkylating agent that triggers replication fork stalling and associated DNA damage signaling ^7,91^, and monitored Mek1 activation status. Despite the rapid induction of DNA damage (as judged by increased phosphorylation of Serine 129 on Histone H2A (phospho-Histone H2A-S129 ^92–94^), expression of *GFP-MEK1* did not lead to an observable effect on phospho-Histone H3-T11 status (Mek1 activity). Thus, expression of GFP-Mek1 did lead to activation of Mek1, even in the presence of upstream DNA damage-induced signaling. We next compared effects on Histone H3-T11 in cells that contain *pGAL1::GFP-MEK1* with cells expressing *pGAL1::GST-MEK1*. Forced dimerization of Mek1 via a GST-fusion leads to (apparently unregulated) Mek1 activation in meiotic cells ^7,95^. Interestingly, we found that expression of *GST-MEK1* in mitotic cells equally led to a strong increase in phosphorylation of Histone H3-T11, within 4 hours of galactose addition (**Figure 6a**, lane 4). In cells expressing *GST-MEK1*, MMS treatment did not enhance the apparent activation of Mek1 (**Figure 6a**). Together, these data suggest that, whereas GST-Mek1 shows (apparently unregulated) activation, expression of GFP-Mek1 is not sufficient to trigger downstream phospho-activation, even in the presence of DNA damage induced upstream signaling. Our observation of increased phosphorylation of Histone H3-T11 upon expression of GST-Mek1 suggests that, also in mitotic cells, fusion of GST with Mek1 leads to (uncontrolled) Mek1 kinase activation, likely via forced dimerization, and that this activation does not require upstream activation.

**Figure 6.**
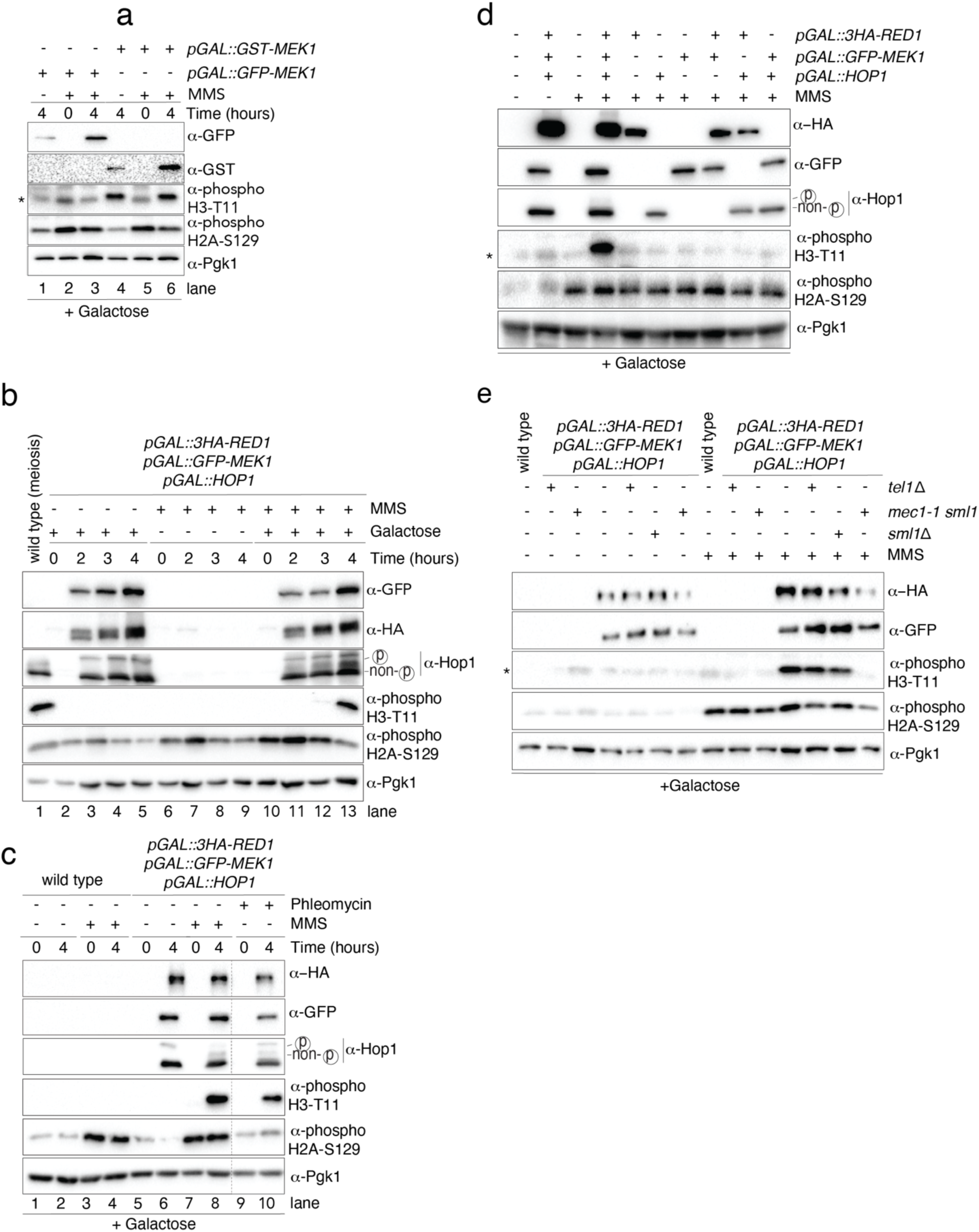
Activation of the Red1-Hop1-Mek1 complex in mitotically-dividing cells. **a**. Analysis of Mek1 activation in mitotically-dividing cells in cells expressing GST-Mek1 (*pGAL::GST-MEK1*; yGV2774) or GFP-Mek1 (*pGAL::GFP-MEK1*; yGV2812). MMS was used to induce DNA damage; cells were treated with MMS for 4.5 hours. 4 hours of galactose-based induction are indicated. See also Supplementary Figure 1 for details on growth and drug treatment conditions. ⍺-phospho-H2A-S129 was used to detect Mec1/Tel1-dependent activation, ⍺-phospho Histone H3-T11 was used to detect Mek1 activation. ⍺-GST and ⍺-GFP was used to detect Mek1. Pgk1 was probed as loading control. * indicates background signal for ⍺-phospho Histone H3-T11 (see main text). **b**. Analysis of Mek1 activation in cells expressing Red1, Hop1 and Mek1 (yGV4806). MMS was used to induce DNA damage; cells were treated with MMS for 4.5 hours. Hours of galactose-based induction are indicated. ⍺-phospho-H2A-S129 was used to detect Mec1/Tel1-dependent activation, ⍺-phospho-Histone H3-T11 was used to detect Mek1 activation. ⍺-GFP was used to detect Mek1, ⍺-Hop1 was used to detect Hop1 (note also the slower migrating band of Hop1, indicating phosphorylation-mediated gel retardation), and ⍺-HA was used to detect Red1. Pgk1 was probed as loading control. A control sample from meiotic wild type cells (yGV49), 4 hours into the meiotic program was used. **c**. Analysis of Mek1 activation in wild type cells (yGV104) and cells expressing Red1, Hop1 and Mek1 (yGV4806). MMS or Phleomycin was used to generate DNA damage. Hours of galactose-based induction are indicated. ⍺-phospho-H2A-S129 was used to detect Mec1/Tel1-dependent activation, ⍺-phospho-Histone H3-T11 was used to detect Mek1 activation. ⍺-GFP was used to detect Mek1, ⍺-Hop1 was used to detect Hop1 (note also the slower migrating band of Hop1, indicating phosphorylation-mediated gel retardation), and ⍺-HA was used to detect Red1. Pgk1 was probed as loading control. **d**. Analysis of Mek1 activation in cells expressing different combinations of RHM complex subunits (yGV104, yGV3726, yGV3243, yGV2812, yGV3235, yGV3255, yGV3219 and yGV4806). MMS was used to induce DNA damage; cells were treated with MMS for 4.5 hours. Galactose-based induction was done for 4 hours. ⍺-phospho-H2A-S129 was used to detect Mec1/Tel1-dependent activation, ⍺-phospho-Histone H3-T11 was used to detect Mek1 activation. ⍺-GFP was used to detect Mek1, ⍺-Hop1 was used to detect Hop1 (note also the slower migrating band of Hop1, indicating phosphorylation-mediated gel retardation), and ⍺-HA was used to detect Red1. Pgk1 was probed as loading control. * indicates background signal for ⍺-phospho Histone H3-T11 (see main text). **e**. Analysis of Mek1 activation in cells expressing Red1, Hop1 and Mek1 in wild type, *tel1Δ*, *sml1Δ, and mec1Δ sml1Δ.* MMS was used to induce DNA damage; cells were treated with MMS for 4.5 hours. Galactose-based induction was done for 4 hours. ⍺-phospho-H2A-S129 was used to detect Mec1/Tel1-dependent activation, ⍺-phospho-Histone H3-T11 was used to detect Mek1 activation. ⍺-GFP was used to detect Mek1, ⍺-HA was used to detect Red1. Pgk1 was probed as loading control. Yeast strains used yGV4806, yGV5011, yGV5033, yGV5044. * indicates background signal for ⍺-phospho Histone H3-T11 (see main text).

We next investigated whether expression of RHM complex subunits might accommodate Mek1 activation. The expression of the entire RHM complex did not trigger an increase in Histone H3-Threonine 11 phosphorylation in cells that did not experience DNA damage (**Figure 6b**). The induction of DNA damage in cells that did not express the RHM complex, did not lead to an increase in H3-Threonine 11 phosphorylation, despite an increase in Mec1/Tel1-dependent phosphorylation of Histone H2A-S129. Strikingly, when we combined the generation of DNA damage (through MMS treatment) with expression of the RHM complex, we observed a specific phosphorylation of Histone H3-T11 after 4 hours of induction (**Figure 6b**, lane 13). Thus, in mitotically dividing cells that expressed the RHM complex and that experienced DNA damage (via MMS treatment), Mek1 kinase can be activated (**Figure 6b**). A main downstream target of Spo11-driven, Mec1/Tel1-dependent signaling that drives Mek1 activation in meiosis is Hop1. Hop1 phosphorylation can be monitored by phospho-specific antibodies or a phosphorylation-induced retardation migration in SDS-PAGE electrophoresis ^48–50^. When we treated cells that expressed the RHMc complex and experienced DNA damage, we noted the presence of a slower migrating form of Hop1 (**Figure 6b**, lane 13) that co-occurred only in conditions where Mek1 was activated. This suggests that indeed activation of Mek1 within the RHMc in mitosis occurs through modification of Hop1 ^48–50^. MMS triggers DNA damage signaling via effects on DNA replication ^91^), which is different from the meiotic DNA damage signaling that occurs during a G2-like state through the generation of Spo11-dependent DSBs ^96^. Generating DNA damage via phleomycin (a DNA break-inducing agent) treatment also led to Mek1 activation (**Figure 6c**, compare lane 8 and 10). Thus, when expressed in mitotic cells, the RHM complex can lead to activation of Mek1 kinase, and that this activation depends on the generation of DNA damage.

This system now enables us to dissect the regulation of this process. Querying Mek1 activation in cells expressing different combinations of RHM components (combined with MMS treatment) revealed that Mek1 was only activated when all three RHM complex components were present, as shown in **Figure 6d**. Thus, as in meiosis, the presence of all three RHM complex subunits is required for efficient Mek1 activation in response to upstream DNA damage signaling. We investigated requirements for upstream Mec1/Tel1-dependent DNA damage signaling. We introduced *mec1Δ* (in an *sml1Δ* background, i.e., *mec1Δ sml1Δ ^97^*) and *tel1Δ* into strains expressing the RHM complex, and treated these cells with MMS to induce DNA damage-dependent signaling. Mutation of *MEC1* led to a marked decrease in phospho-Histone H3-T11 signal, whereas under similar treatment conditions, Tel1 did not appear to play a significant role in Mek1 activation (**Figure 6e**). These observations are in line with earlier experiments analyzing the regulation of RHM complex activation (via Hop1 phosphorylation) in meiosis ^48,91,49–51^: also here, Mec1 is mainly responsible for Hop1 phosphorylation (and thus Mek1 activation).

Expressing and activating the RHM complex components in mitosis in principle now allows us to query Mek1 activation mechanisms in ways that are difficult to achieve in meiosis. We first aimed to address potential cell cycle-dependent regulation of Mek1 activation. In meiosis, the RHM complex only becomes active in meiotic G2/prophase, concomitantly with Spo11-dependent DSB formation. Our mitotic activation system allowed us to test if Mek1 could be activated outside of a G2-like cell cycle state. For this, we combined the expression of *RED1*, *HOP1* and *MEK1* with DNA damage induction and cell cycle synchronization in G1 (⍺-factor treatment) or G2/M (nocodazole treatment). For this experiment, we employed phleomycin as a DNA damaging agent, because MMS is not expected to lead to DNA damage outside of S-phase. Indeed, treatment of MMS in synchronized cells did not lead to detectable activation of Mec1/Tel1-based signaling (*i.e*., in G1-arrested cultures, **Supplementary Figure 6a**). Cells arrested in G2/M readily activated Mek1, in a manner similar to the activation seen in asynchronously growing cells (**Figure 7a**). However, we found that cells that were arrested in G1 were unable to trigger activation of Mek1 upon phleomycin treatment, despite the activation of upstream Mec1/Tel1 signaling triggered by phleomycin treatment (**Figure 7a**). These observations suggest that in G1 phase activation of the RHM complex activation. What could be the reason behind this observation? A possibility is that it is inherent to the RHM complex - *i.e.*, that a crucial modification or event (that normally occurs in meiotic G2/prophase) is lacking in this phase of the mitotic cell cycle, which might preclude RHM complex formation and/or Mek1 activation. We investigated if RHM complex formation was affected in cells that were synchronized in G1 phase of the cell cycle (by ⍺-factor). Co-immunoprecipitation experiments revealed that the RHM complex could be formed efficiently, also in G1-arrested cells (**Figure 7b**), thus likely excluding a failure in RHM complex assembly in G1 phase of the cell cycle as an underlying reason for the failure to active Mek1. A second possibility was that upstream signaling upon DNA damage induction is lacking or inefficient. Our data (**Figure 6e**) and that in earlier work ^48,91,49–51^ suggests that RHM complex activation relies on an efficient Mec1 function. Mec1 activation depends on ssDNA resection, and ssDNA resection at DNA breaks is limited in G1 phase ^98–100^. Reduced activation of Mec1 – due to diminished DNA end resection in G1 – might conceivably thus underlie the failure to activate the RHM complex in G1 phase of mitotic cells.

**Figure 7.**
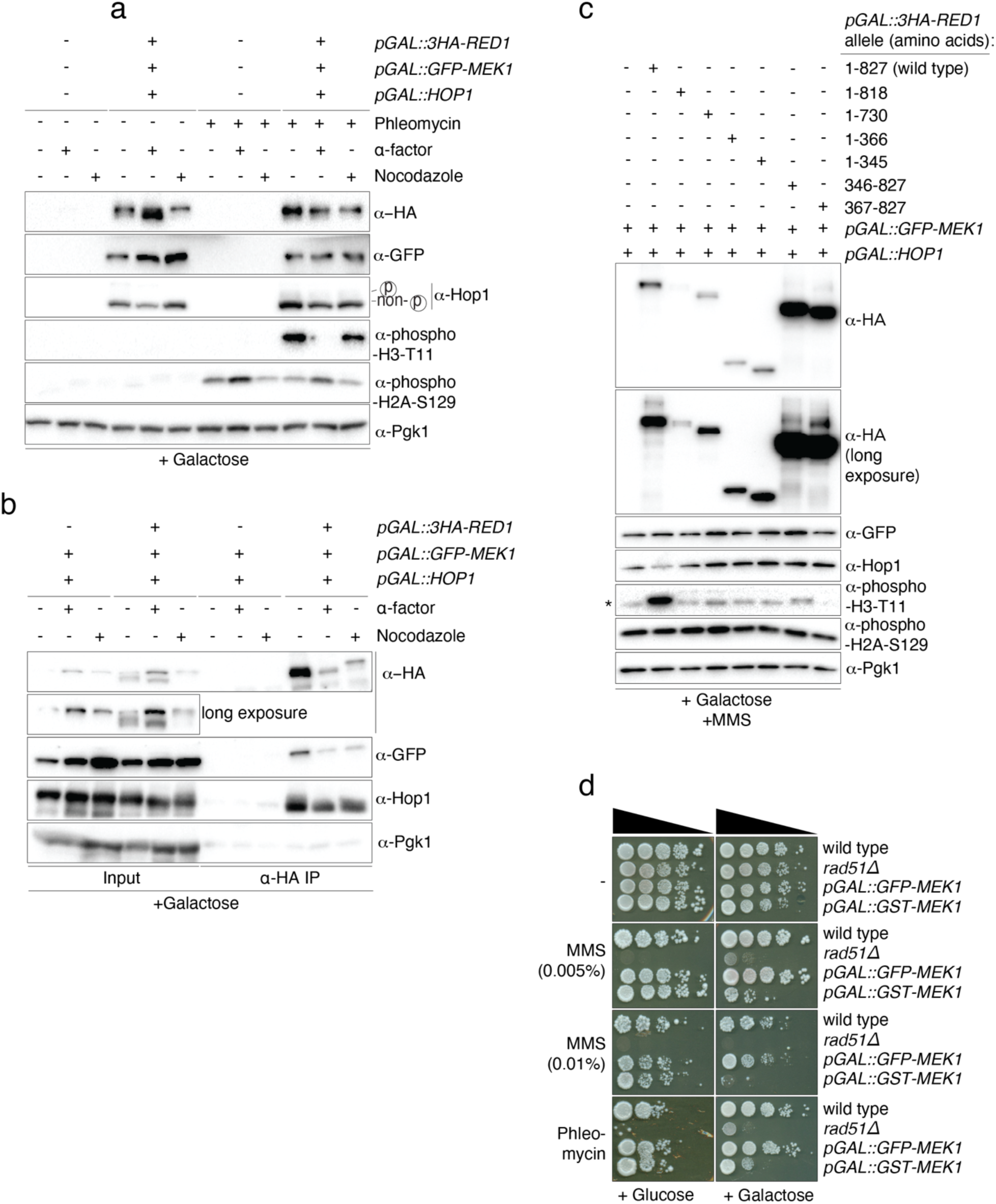
Red1-Hop1-Mek1 complex assembly and activation during the cell cycle and in the presence of Red1 truncation alleles. **a.** Analysis of Mek1 activation in wild type or Red1, Hop1 and Mek1 expressing cells under different cell cycle conditions. Yeast strains used are yG104 and yGV480. Cells were arrested in G1-phase by the addition of ⍺-factor, and in mitosis by addition of nocodazole (see Material and Methods and Supplementary Figure 1 for details). Phleomycin was used to induce DNA damage. Galactose-based induction was for 4 hours. ⍺-phospho-H2A-S129 was used to detect Mec1/Tel1-dependent activation, ⍺-phospho-Histone H3-T11 was used to detect Mek1 activation. ⍺-GFP was used to detect Mek1, ⍺-Hop1 was used to detect Hop1 (note also the slower migrating band of Hop1, indicating phosphorylation-mediated gel retardation), and ⍺-HA was used to detect Red1. Pgk1 was probed as loading control. **b**. Co-immunoprecipitation (co-IP) between Red1 and Hop1/Mek1 under different cell cycle conditions. Cells were arrested in G1-phase by the addition of ⍺-factor, and in mitosis by addition of nocodazole (see Material and Methods for details). Red1 was immunoprecipitated via ⍺-HA pulldown. ⍺-HA was used to detect Red1, ⍺-GFP was used to detect Mek1, and ⍺-Hop1 was used to detect Hop1. Pgk1 was probed as loading control. * indicates background signal. Samples were taken after 4 hours induction with galactose. Yeast strains used: yGV3219 and yGV4806. **c**. Analysis of Mek1 activation in mitotically dividing cells in cells expressing Hop1 and Mek1, combined with different Red1 truncations. MMS was used to induce DNA damage; cells were treated with MMS for 4.5 hours. Galactose-based induction was for 4 hours. ⍺-phospho-H2A-S129 was used to detect Mec1/Tel1-dependent activation, ⍺-phospho-Histone H3-T11 was used to detect Mek1 activation. ⍺-GFP was used to detect Mek1, ⍺-Hop1 was used to detect Hop1, ⍺-HA was used to detect Red1. Pgk1 was probed as loading control. Strains used were: yGV3219, yGV4806, yGV4393, yGV4395, yGV4397, yGV4400, yGV4207, and yGV4402. * indicates background signal for ⍺-phospho Histone H3-T11 (see main text). **d**. Serial dilution (10-fold) spotting of yeast strains, on Glucose-or Galactose-containing solid medium, containing MMS (0,005% and 0,01%) or Phleomycin. Strains used were: yGV104, yGV2774, yGV2812, yGV3753.

In addition to being essential in establishing the RHM complex, Red1 and Hop1 also play important roles in establishing the meiotic chromosome loop-axis organization which is key in mediating Spo11 activation ^20,4,11,12,24–26^. Indeed, mutations in Red1 and/or Hop1 lead to diminished Spo11-driven DNA break formation in meiosis. Since Spo11-dependent DNA break formation is required for Mec1/Tel1-dependent Mek1 activation, any effects on meiotic DNA break activity complicate the interpretation effects of Red1 or Hop1 mutations on Mek1 activation. Our mitotic RHM complex expression and activation system enables us to in principle uncouple the activation of Mek1 from DNA break-dependent signaling. We therefore used our system to investigate how different truncated Red1 alleles (see earlier and **Figure 2c**) affected Mek1 activation upon DNA damage induction. We expressed the described Red1 truncation alleles – in combination with Hop1 and Mek1 – and monitored the activation of Mek1 upon DNA damage induction (**Figure 7c**). Under these experimental conditions, only the expression of wild type (*i.e*., full length Red1^1-827^) led to a detectable activation of Mek1 (as judged by phosphorylation of Histone H3-T11). Thus, the integrity of the RHM complex is crucial to enable Mek1 activation. For example, we observed that the presence of the COOH-terminal region of Red1 is needed for Mek1 activation (**Figure 7c**, Red1^1-366^), even though this truncated version of Red1 was able to interact with Hop1 as well as Mek1 (**Figure 2d)**. Expression of a truncated version of Red1 that expresses the COOH-terminal region of Red1 but lacks its NH_2_-terminal ARM/PH domain (Red1^346-827^) was equally not able to activate Mek1. In agreement with the idea that the NH_2_-terminal region is however not sufficient for activation was the observation that the removal of the COOH-terminal coiled-coil domain (amino acids 737-827; responsible for tetramerization and filament formation of Red1 ^12,13,20^ was associated with a lack of Mek1 activation upon expression of this fragment (Figure 7c, Red1^1-730^). Hop1 and Mek1 were able to interact with this truncated Red1 protein (**Figure 2d**). In combination with previous work ^12,13,20^, this suggests that in our mitotic system, tetramerization and/or filament formation of the RHM complex was needed for efficient Mek1 activation. Further truncation revealed that removing the most COOH-terminal 9 amino acids of Red1 (*i.e.*, 819-827) was also associated with a failure in Mek1 activation (**Figure 7c**, Red1^1-818^), whereas this Red1 protein efficiently interacted with Hop1 and Mek1 (**Figure 2d**). Removing this amino acid stretch is associated with defects in meiotic G2/prophase and was suggested to specifically interfere with the filamentous assembly of Red1 (and by interference RHM complexes)^13,96^. Thus, also in our system, activation of Mek1 (driven by exogenous DNA damage) appears to require filament formation of the RHM complex, an activity that is encoded in the extreme COOH-terminus or Red1 ^12,13,20^.

We finally evaluated functional consequences of RHMc/Mek1 kinase activation on (mitotic) DNA repair. The current model of RHMc-mediated DNA repair modulation posits that active Mek1 is localized, and as such establishes a localized HR inhibition ^63^, most prominently mediated through phosphorylation of the Rad51-accessory factor Rad54 ^6,7,56,57,61,62^. Localized inhibition of Rad51-dependent HR could, with the help of additional meiosis-specific events, eventually lead to IH-based repair. The abovementioned growth defect associated with the expression of RHMc (*i.e.*, see **Figure 1f**) precluded us from using these strains to explore effects of RHMc activation on growth in DNA damaging conditions. We instead explored effects of activation Mek1 via the expression of the constitutively active GST-Mek1 (**Figure 5a**). As shown in **Figure 7a**, expression of GST-Mek1 (in contrast to the expression of GFP-Mek1) led to a growth defect specifically in cells that experienced DNA damage (either by MMS or phleomycin treatment). The observed effects were similar to effects seen in *rad51Δ* cells (**Figure 7a**), and in mitotic cells that express a Mek1-driven phosphomimic *RAD54* allele ^7^. This suggests that activating Mek1 in mitotic cells is sufficient to establish certain characteristics that typify meiosis-specific DNA repair. In total, our work presents the development of two heterologous systems to interrogate RHMc structure, assembly and function. We use this to reveal organizational principles that govern the assembly and activation of this central regulator of meiosis-specific HR-based repair. Our experiments in mitotic cells indeed point to a role for the RHMc in inhibiting ‘mitotic’ intersister-based DNA repair.

## Discussion

Here, we study the establishment and activation of the budding yeast Red1-Hop1-Mek1 complex. We express components of the trimeric, meiosis-specific Red1, Hop1 and Mek1 complex in mitotically dividing yeast cells. We show that this complex can self-assemble when expressed outside of its natural, physiological environment, *i.e.,* in mitotically dividing cells (**Figure 1 and 2**). Thus, the RHM complex does not require additional meiosis-specific factors or modifications for its assembly. This is somewhat surprising, since earlier work in meiosis has suggested that the incorporation of Mek1 into the Red1-Hop1 complex requires upstream input in the form of (mostly) Mec1-dependent phosphorylation events (most notably on Hop1) ^48^. We cannot currently exclude the possibility that Mec1-dependent phosphorylation during DNA replication is sufficient to promote an interaction-competent state. We note that in original work on the RHMc, a yeast-2-hybrid experiment detected an autonomous association of Mek1 with Red1^19^. Regardless, our observations in mitotically-dividing cells are supported by *in vitro* reconstitution efforts coupled to *in silico* modeling, clearly suggesting that the association between Red1 and Hop1 involves more than the already described ‘closure motif’-HORMA binding ^13^. Furthermore, it suggests the presence of (a) direct binding interface(s) between Mek1 and Red1.

Our *in vivo* and *in vitro* data confirms the idea that the capture of the CM within Red1 (Red1^340-362^) by the HORMA domain of Hop1 is a key step in RHMc assembly ^13,19,12^ (**Figure 2a** and **5**). Based on the known biochemical and structural behavior of HORMA domains (^101–103^), we presume that this association involves a ‘closed’ Hop1^HORMA^ topology, in which a CM-HORMA ‘seat belt’ binding captures Red1. In addition to this expected mode of interaction, our data points to additional binding interfaces between Red1 and Hop1 (**Figure 5 and Supplementary Figure 5a**). First, our *in vivo* co-IPs suggest that the NH_2_-terminal region immediately upstream of the CM (Red1^1-345^) contributes to binding (**Figure 2e**, compare lane 5 and 6). Second, in silico structural modeling with AlphaFold2Multimer points to the existence of interaction interfaces within the ARML and PH domains that make up the NH_2_-terminal region of Red1 (**Figure 5a and b**). Indeed, our subsequent *in vitro* reconstitution enforces this idea (Figure 5). In total, we suggest that upon capture of the CM by the HORMA domain of Hop1, the ‘closed’ HORMA can establish several interactions with the nearby ARML-PH assembly (**Figure 5h**). This interaction could stabilize the association of Hop1 with Red1, potentially by preventing a simple ‘slide-off’ disengagement of CM peptide form within Hop1^HORMA^, as has been demonstrated in case of CM-mediated association of Cdc20 with the HORMA protein Mad2 ^80^. Alternatively, it might trigger conformational changes within the NH_2_ terminus of Red1, to accommodate other binding partners. A speculative consequence of such a rearrangement might be the accommodation of Mek1 interaction by either Red1 or Hop1, or the recruitment of other Hop1 binders, such as Mer2, ensuring that Mer2 preferentially associates with axis-localized Hop1 ^15^. Finally, this constellation might influence Red1-Hop1 disassembly dynamics. The AAA+ ATPase Pch2/TRIP13 catalyzes the transition of Hop1’s HORMA domain from ‘closed’ to ‘open’, as such releasing CM-mediated associations ^101,104^. Interaction of the Hop1^HORMA^ with the Red1^ARML-PH^ domains could conceivably impact this reaction, either protecting Hop1 from Pch2/TRIP13 or promoting its enzymatic removal from the axis. Our data further points to the fact that Mek1 can directly associate with Red1 (**Figure 3** and **4**). Interestingly, our modeling efforts suggest a possible mode of interaction between the kinase domain of Mek1 and the ARML domain of Red1 that structurally resembles the described interaction between the human STK24 kinase and its scaffolding activator MO25b ^82,83-84^ (**Figure 3**). This prediction agrees with our subsequent biochemical experiments (**Figure 3**). An important corollary from our modeling is that to accommodate the observation that a ‘STK24-MO25-like’ organization would necessitate significant spatial movements of the adjacent Red1^PH^ domain to make binding to the Red1^ARML^ domain with Mek1 possible (**Figure 3e**). One possibility is that the Red1^PH^ domain binds directly to, for example, a phosphorylated protein, consistent with the typical role of PH domains. The similarities predicted by these structures highlight that the assembly of a functional RHMc like involves several intermolecular associations between subunits. In addition, it proposes the possibility that, in addition to being driven by phospho-regulated dimerization of Hop1 ^7,48,49^, Mek1 kinase activation might be regulated via associations with Red1. Clearly, future work should be aimed at addressing these important questions.

Using our mitotic expression system, we demonstrate that the RHM complex can be integrated into mitotic DNA damage-induced kinase signaling, which eventually leads to the activation of Mek1 kinase (**Figure 6** and **7**). Thus, the RHM complex is activated in a non-physiological environment. Our work lays the groundwork for further exploration of potential effects of RHM complex activation beyond meiosis on DNA repair dynamics and decisions aimed at testing current models of interhomolog bias establishment (see also below). The initial experiments we describe here provide a proof-of-concept showcasing the feasibility of our experimental approach. We use our system to investigate the requirements for Mek1 activation. We find that all three components of the RHM complex are required for Mek1 activity upon treatment with DNA damaging agents – unless Mek1 is artificially dimerized through GST-fusion, as was also observed in meiotic cells ^7,95^ (**Figure 6**). Mitotic activation of the RHM complex in response to DNA damaging conditions relies to a large extent on Mec1 function, mirroring observations made in meiotic cells ^48–51^. Furthermore, we found that the activation of Mek1 by upstream Mec1-dependent signaling is impaired in G1-phase of the cell cycle. Mec1 activity is dependent on ssDNA generation at DNA breaks ^105,106^, and DNA end resection is inefficient in G1 phase due to low Cdk1 activity ^98–100^. We suggest that efficient activation of the RHM complex (in mitosis) requires efficient DNA end resection-mediated Mec1 activity. We probe the dependency of RHM complex formation and activation on Red1 structural domains. Using this approach, we find that although RHM complex formation can be established in several Red1 mutants lacking certain structural and functional domains (**Figure 2d and e**), the activation of Mek1 appears to require full length Red1 protein (**Figure 4c**). We suggest that the highly controlled assembly and activation of the RHM complex is responsive and reliant on several (upstream) inputs, such as phosphorylation events, chromatin association, filamentous assembly, which eventually leads to the promotion of Mek1 activation. For example, our data indicate that the NH_2_-terminal ARM/PH domain of Red1 ^66,77,78^ is required for the activation of Mek1. In addition to the discovered association with Hop1 and Mek1 (see above), this domain might mediate the association between Red1 and meiotic (Rec8)-containing cohesin ^13,17^. Thus, chromosomal association might be key to enabling Mek1 activation and subsequent downstream phosphorylation. It will be key in the future to establish whether the RHM complex is recruited to mitotic chromosomes, and if so, where it is recruited and whether association is dependent on the NH_2_-terminal ARML/PH domain of Red1 (and on cohesin).

We were so far unable to reconstitute a Mek1-Red1-Hop1 complex using *in vitro* approaches. This might indeed indicate that Mek1-assocation is influenced by additional cellular factors not included in our purified systems. *In silico* modeling of the interactions between Red1^AMRL-PH^, Hop1 and Mek1 suggest that extensive structural rearrangements (as for example a re-organization of Red1^PH^, see also **Figure 3**) are needed to accommodate a 1:1:1 assembly (**Supplementary Figure 7a**). Thus, it will be a central future goal to understand the stoichiometries and minimal assembly requirements between these three factors. It should be noted that genetic evidence points to a Hop1-dependent dimerization step that is needed for Mek1 activation ^7^, whereas Red1 (through its COOH-terminal coiled-coil domain) can also establish multimeric assemblies ^7,13^. Indeed, our experimental analysis further suggested that the extreme COOH-terminal region of Red1, including its filament forming domain, is needed to endow the RHM complex with Mek1-activating functionality (**Figure 7**). In addition to the role of a possible chromatin-interacting domain (in the NH_2_-terminus of Red1), the proper assembly of large-scale filaments might enable Mek1 activity, potentially via enhancing (chromatin-associated) substrate availability. Earlier work has shown that filament formation of Red1 is needed for the proper assembly of the meiotic chromosome axis, and for successful meiosis ^13,20,79,107^. Higher-order organization of the RHM complex might thus also more directly contribute to efficient activation of Mek1 kinase activity. How this organization contributes to Mek1 activity, and whether it involves facilitating substrate availability or interaction, should be a key future question. Answering this question likely requires combining *in vivo* analysis with *in vitro* biochemical reconstitution of functional RHM complexes.

Expression of Red1, a large protein that can form filamentous assemblies ^13,20^, is associated with growth defects likely stemming from delayed mitotic progression (**Figure 1j, Supplementary Figure 2a, Figure 4f)**), in agreement with earlier work ^76^. Cell cycle delay triggered by Red1 expression is exacerbated under conditions where Hop1, Mek1 or Hop1/Mek1 are co-expressed (**Figure 1f and g**). The nature of these effects is unknown, but it is conceivable that these are caused by the establishment of large protein assemblies/filaments made up of Red1, and/or Hop1, potentially on mitotic chromosomes. A structure-based mutagenesis analysis with regards to the observed cell cycle effects revealed that indeed the COOH-terminal region of Red1 (responsible for the Red1 multimerization and filament formation ^13,20^) was involved and seemed sufficient to trigger cell cycle effects (**Figure 4f**). We note that in addition to driving Red1 filaments, this region was shown to interact with the 9-1-1 DNA damage complex components Mec3 and Ddc2 ^79^ and SUMO (Smt3) chains ^13,20,79,107^. Currently, we cannot distinguish whether interaction with any of these factors is associated with the cell cycle arrest we observed, or whether it is indeed a consequence of filament assembly. Further work should be aimed at addressing these questions.

The availability of experimental tools and the characteristics of mitotic cells will allow us to query the role of the RHM complex – and potential sufficiency thereof – in establishing alterations in DNA repair. In mitosis, HR-based repair is primarily executed using repair templates that are present on sister chromatids ^3^, whereas the RHM complex is crucial in promoting interhomolog-biased repair during meiotic HR-based DSB repair. How the RHM complex – and thus the Mek1 kinase – achieves this sister-to-homolog switch during DBS repair is incompletely understood. Several phospho-substrates of Mek1 have been characterized, and a shared functional consequence of Mek1 function is an inhibitory effect on Rad51-mediated DNA repair ^6,7,56,57,61,62^. Most prominently, Mek1 phosphorylates the Rad51-accessory factor Rad54 ^6,7,56,57,61,62^. This reduces the ability of Rad51 to promote homologous recombination ^6,7,56,57,61,62^. The current model posits that a sister chromatid-restricted Mek1-activity establishes a localized zone on meiotic chromosomes that is not permissive to HR-based DNA repair ^63,64^. Escape from this localized activity – with the goal of allowing DSB repair – could only be achieved by distant repair events, which can be found on homologous chromosomes. As such, local Mek1-driven HR inhibition would essentially encourage homology-based repair through local inhibition of DSB repair. We can now in principle explore this model using our mitotic RHM-activation system. It will allow us to ask a myriad of questions, including but not limited to: *i)* is sister-based HR repair inhibited by RHM complex activation in mitosis? *ii)* Is this associated with reduced Rad51 function, and *iii)* a possible increase in homology-directed repair? *iv)* What are the potential effects of this kind of repair on cell cycle progression and genome stability in mitosis? We tested the first question, with regards to RHMc function. Expression of GST-Mek1 – a constitutively active Mek1^7,95^ – in mitotic cells leads to sensitivity to DNA damaging conditions, in a manner that is comparable to cells that lack Rad51 (**Figure 7d**). These experiments suggest that the activation of Mek1 thus is sufficient to inhibit HR, also in mitotic cells. Hence, these findings agree with the proposed model by which Mek1 can lead to a local inhibition of Rad51-dependent repair ^63^. We performed these experiments in haploid cells (*i.e.,* in cells that do not contain homologous chromosomes as alternative repair templates). It will be interesting to explore functional consequences of Mek1 activity in diploid cells, especially in combination with the utilization of directed DNA damage-inducing systems and physical assays to monitor HR dynamics (*e.g.*, ^3^). Several other meiotic factors, such as Rec8 and Hed1, contribute to interhomolog-directed HR bias in meiosis ^7,56,75,108,109^. To establish a ‘more complete’ switch to meiosis-like HR repair, it might thus be required to combine our system with the expression of additional factors.

The human genome encodes functional homologs of RHM components (*e.g.,* HORMAD1/HORMAD2 – Hop1 homologs, and SYCP2/SCYP3 – Red1 homologs) (reviewed in ^66,101^), and mutations in SYCP2 or HORMAD1 lead to infertility ^63,110,111^. These factors likely influence interhomolog DNA repair bias. We finally note that human RHM complex components are frequently aberrantly re-expressed in cancer and alter DNA repair pathways under pathological conditions (*e.g.,^66,101,112,113^*^-114,115^). Our work establishes a conceptual framework and tractable system to query functionality of human meiosis-specific DNA repair factors and consequences of aberrant expression of these factors on genome maintenance during tumorigenesis.

## Methods

### Yeast strain construction and genetics

Genotypes of budding yeast (*Saccharomyces cerevisiae*) strains are listed in **Supplementary Table 1**. For construction of galactose-inducible expression alleles, *pGAL1*, *pGAL1::GFP, pGAL1::GST* or *pGAL1::3HA* constructs were integrated directly upstream of the start codons of *HOP1*, *MEK1* or *RED1* using PCR-based gene targeting ^116^. *RED1* NH_2-_-terminal truncations were generated by integration of *pGAL1::3HA* at desired codon sites. COOH-terminal truncations were made by introduction of a STOP codon (via integration of a *TRP1* marker including a STOP codon ^116^) in a *KANMX6::pGAL1::3HA-RED1* containing strain. Integration of constructs was confirmed by PCR. Subsequent strains were generated by yeast genetics and tetrad dissection.

### Growth conditions, galactose-based expression induction and drug treatments

Strains were grown on YP-glycerol plates, transferred to YP-glucose (YPD) plates and grown overnight. Cultures were grown in YP-raffinose/glucose (YPRG) media (2.4% Raffinose, 0.12% Glucose) overnight at 30°C till saturation. In the morning, cultures were diluted to a final OD_600_ of 0.48 and further grown for 3.5 hours in YP-RG before adding 2% Galactose. Samples were collected at indicated time points. For DNA damage drug treatment, 0.01% (v/v) methyl methanesulfonate (MMS, Sigma-Aldrich) was added 30 minutes prior to galactose induction. Alternatively, phleomycin (Sigma-Aldrich) treatment was performed by adding 50 𝜇g/ml (30 minutes prior to galactose addition). Phleomycin was re-added 2 hours after galactose-based induction. For enrichment of cells in G1, secondary cultures were grown for 2.5 hours followed by the addition of α-factor (166 𝜇g/ml; home-made). A second dose of α-factor (50 𝜇g/ml) was added after 1 hour of galactose induction. For nocodazole-induced G2/M arrest, secondary cultures were treated with nocodazole (15 𝜇g/ml; Sigma-Aldrich) 3 hours after starting secondary cultures. Nocodazole (15 𝜇g/ml) was re-added 3 hours after adding galactose. For meiotic time courses (Figure 1D and 3b), cells were treated and cultured as described in detail in ^63,117^.

### Yeast viability assays

Cells were inoculated into liquid YP-RG media overnight at 30°C. The next day, cells were diluted to OD_600_ 1 in H_2_O. 5 μl of 10-fold dilutions were spotted on YP-glucose or YP-galactose plates. Growth at 30°C was monitored for the following 2–4 days.

### Cell cycle analysis

Flow cytometry was used to monitor cell cycle status. Briefly, 150 µL of yeast cultures were fixed (2 hours at 4 °C) in 70% EtOH. Fixed cells were pelleted and incubated overnight at 50 °C in 500 µL Sodium Citrate (50 mM) with 0.7 mL RNase A (30 mg/ml; Sigma-Aldrich). Cell suspensions were treated with proteinase K (20 mg/ml; VWR) for 2 hours at 50 °C. DNA was stained by addition of 500 µL of Sodium Citrate (50 mM) containing 0.2 µL SYTOX-Green (Life Technologies). Cells were disrupted by brief sonication, and DNA content was measured using a BD AccuriTM C6 (BD Biosciences) flow cytometer. Samples were taken at the indicated time points, and 10,000 events were counted. Data was analyzed using FlowJo (FlowJo LLC).

### SDS-PAGE and western blotting

Samples were harvested at indicated time points by harvesting the equivalent of 5 ml OD_600_ 1.9 from the cultures. Cell pellets were precipitated with 5 ml 5% TCA and washed with acetone. Pellets were dried overnight and were resuspended in 200 ml of protein breakage buffer (10mM Tris (pH 7.5), 1 mM EDTA, 2.75 mM DTT). ∼0.3g of acid-washed glass beads was added, and cell breakage was performed by using a FastPrep-24 (MP Biomedicals). Samples were diluted by adding 100 μl of protein breakage buffer and 150 μl of 3x SDS loading buffer. To observe histone modifications, 15% SDS-PAGE gels were used. 15% Gels were run at 70 Volt for 100 minutes. Protein transfer was done on PVDF membranes (phospho-histone detection) or on nitrocellulose membranes (other proteins). Primary antibodies were used as follows: α-HA (Biolegend 901502; 1:500), α-Pgk1 (Thermo Fisher; 1:1000), α-GFP (home-made; 1:5000), α-Hop1 (home-made; 1:10,000, see ^118^), α-phospho-Histone H3-Threonine 11 (EMD Millipore 05-789, 1:1500), and α-phospho-Histone H2A-Serine 129 (Abcam 181447, 1:500 in 4% BSA/TBS-Tween). All antibody incubations were in 5% milk/PBS-Tween, unless stated otherwise.

### Co-immunoprecipitation

Secondary cultures were grown for 3.5 hours followed by 2% galactose addition for 4 hrs. The equivalent of 50 ml of cultures (OD_600_ 1.9) were spun down at 3000 rpm for 3 minutes. Samples were washed with cold H_2_O and snap frozen. 300 μl of ice-cold IP buffer (50 mM Tris–HCl pH 7.5/150 mM NaCl/1% Triton X-100/1 mM EDTA pH 8.0, with a cocktail of protease inhibitors freshly added) and acid-washed glass beads were added. Cells were broken in a FastPrep-24 disruptor (MP Biomedicals) by two 50-s cycles (speed 6). Cell lysate was spun 30 seconds at 500 rpm, and the supernatant was transferred to a 15 mL falcon tube, followed by sonication for 25 cycles (30 seconds on/30 seconds off, high power range) on a Bioruptor-Plus sonication device (Diagenode), and spun down 30 minutes at maximum speed. Supernatant (total of 500 μl) was transferred to a new microcentrifuge tube, and 10% of the supernatant (50 μl) was collected as input. For ⍺-HA IPs, 1 μl of antibody (⍺-HA; BioLegend) was added to the lysate and rotated for 3 h. Lysates were incubated with 30 μl of Dynabeads protein G (Invitrogen, Thermo Fisher Scientific) overnight at 4°C. For ⍺-GFP-based IPs, 1 μl ⍺-GFP (home-made) were used, in combination with 30 μl of Dynabeads protein G. After incubation, beads were washed four times with 500 μl of ice-cold IP buffer. For the final wash, beads were transferred to a new microcentrifuge tube and resuspended in 40 μl IP buffer. 20 μl of SDS-loading buffer was added to samples and samples were incubated for 5 minutes at 95° C. Inputs were treated as follows: supernatant was precipitated with 10% TCA (*i.e.* 5 μl), and samples were incubated on ice for 30 minutes. Pellets were collected by centrifugation (1 minute at maximum speed), and washed with ice-cold acetone. After centrifugation and removal of supernatant, precipitations were dried on ice, resuspended in TCA-resuspension buffer (50 mM Tris–HCl 7.5/6 M urea), and incubated on ice for 30 minutes. Precipitates were dissolved by careful pipetting and vortexing. 10 μl of SDS-loading buffer was added, and samples were incubated for 5 minutes at 95°C. IP and input samples were analyzed by SDS-PAGE followed by western blotting.

### Protein purification

#### Hop1

Full-length Hop1 constructs (WT, K593A) were produced as 3C HRV cleavable N-terminal Twin-StrepII tag fusion proteins in BL21 STAR *E. coli* cells. Cell cultures were grown at 37°C shaking at 150 rpm until an OD of 0.6 was achieved. Protein expression was induced by the addition of 250 µM IPTG, and expression continued at 18°C for 18 hr. Harvested cells were resuspended in lysis buffer (50 mM HEPES pH 7.5, 300 mM NaCl, 7.5% glycerol, 1 mM MgCl_2_, 0.1% Triton-X) and lysed by sonication in the presence of SERVA protease (50 µg/mL), AEBSF protease (50 µg/mL) and DNAseI (20 µg/mL) and cleared through ultracentrifugation at 35,000 rpm for 45 minutes at 4°C (Beckman Coulter). Cleared lysate was applied to a 5 mL Strep-Tactin™XT Superflow™ column (IBA Lifesciences) pre-equilibrated in lysis buffer. A 2 column volume (CV) ATP wash (lysis buffer supplemented with 1 mM ATP) and high salt (50 mM HEPES, 800 mM NaCl, 7.5% glycerol) was performed before extensive washing in lysis buffer (15 CV). The bound protein was eluted with lysis buffer containing 50 mM biotin and loaded on a HiTrap Heparin HP column (GE Healthcare) and subsequently eluted with increasing salt gradient to 1 M NaCl. Eluted strep-Hop1 constructs were concentrated on a 30 kDa MWCO Amicon concentrator and loaded on a Superdex 200 10/300 gel filtration column (GE Healthcare) pre-equilibrated in SEC buffer (50 mM HEPES pH 7.5, 300 mM NaCl, 7.5% glycerol, 1 mM MgCl_2_, 1 mM TCEP). Fractions containing purified strep-Hop1 were concentrated, flash frozen and stored at -70°C.

Hop1 HORMA domain was produced as a 3C HRV cleavable N-terminal Twin-StrepII tag fusion protein in BL21 STAR *E. coli* cells. Cell cultures were grown at 37°C shaking at 150 rpm until an OD of 0.6 was achieved. Protein expression was induced by the addition of 250 µM IPTG, and expression continued at 18°C for 18 hr. Harvested cells were resuspended in lysis buffer (20 mM Tris-HCL pH 8.5, 250 mM NaCl, 7.5% glycerol, 1 mM MgCl_2_, 0.1% Triton-X, 25 mM arginine and glutamic acid), lysed by sonication in the presence of SERVA protease (50 µg/mL), AEBSF protease (50 µg/mL) and DNAseI (20 µg/mL) and cleared through ultracentrifugation at 35,000 rpm for 45 minutes at 4°C (Beckman Coulter). Cleared lysate was applied to a 5 mL Strep-Tactin™XT Superflow™ column (IBA Lifesciences) pre-equilibrated in lysis buffer. A 2 CV ATP wash (lysis buffer supplemented with 1 mM ATP) and high salt wash (20 mM Tris-HCL pH 8.5, 800 mM NaCl, 7.5% glycerol, 1 mM MgCl_2_, 25 mM arginine and glutamic acid) was performed before extensive washing in lysis buffer (15 CV).The bound protein was eluted with a lysis buffer containing 50 mM biotin and loaded on a HiTrap Q column (GE Healthcare) and subsequently eluted with increasing salt gradient to 1 M NaCl. Fractions containing purified strep-Hop1 HORMA domain were concentrated on a 10 kDa MWCO Amicon concentrator, flash frozen and stored at -70°C.

#### Mek1

Mek1 was produced as a 3C-HRV cleavable C-terminal Twin-StrepII tag fusion protein in insect cells. Expression plasmids were used to generate bacmids via the EmBacY cell line, and subsequently transfected into SF9 cells using FuGene HD (Promega). Baculovirus was generated through three rounds of amplification in SF9 cells grown in Sf-900 III media (ThermoFisher), shaking 150 RPM at 27°C. For protein expression, Hi5 cells were infected with the amplified Mek1-strep baculovirus at a ratio of 1:100 (v/v ratio), and cells were cultured for 72 hrs post infection. Harvested cells were washed with PBS, resuspended in lysis buffer (50 mM HEPES pH 7.0, 300 mM NaCl, 7.5% glycerol, 1 mM MgCl_2_, 0.1% Triton-X), lysed by sonication in the presence of SERVA protease (50 µg/mL), AEBSF protease (50 µg/mL) and DNAseI (20 µg/mL) and cleared through ultracentrifugation at 40,000 rpm for 45 minutes at 4°C (Beckman Coulter). Cleared lysate was applied to a 5 mL Strep-Tactin™XT Superflow™ column (IBA Lifesciences) pre-equilibrated in lysis buffer. A 2 CV ATP wash (lysis buffer supplemented with 1 mM ATP) and high salt wash (50 mM HEPES, 800 mM NaCl, 7.5% glycerol, 1 mM MgCl_2_) was performed before extensive washing in lysis buffer (15 CV). The bound protein was eluted with a lysis buffer containing 50 mM biotin and concentrated on a 30 kDa MWCO Amicon concentrator and loaded on Superdex 200 10/300 pre-equilibrated in SEC buffer. Fractions containing purified Mek1-strep were concentrated, flash frozen and stored at -70°C.

#### Red1

Red1^I743R^ was produced as a 3C HRV cleavable C-terminal MBP fusion protein in insect cells. Expression plasmids were used to generate bacmids via the EmBacY cell line, and subsequently transfected into SF9 cells using FuGene HD (Promega). Baculovirus was generated through three rounds of amplification in SF9 cells grown in Sf-900 III media (ThermoFisher), shaking 150 rpm at 27°C. For protein expression, Hi5 cells were infected with the amplified Red1-MBP baculovirus at a ratio of 1:100 (v/v ratio), and cells were cultured for 72 hrs post infection. Harvested cells were washed with PBS, resuspended in lysis buffer (50 mM HEPES pH 7.5, 300 mM NaCl, 10% glycerol, 1 mM MgCl_2_, 0.1% Triton-X, 1 mM EDTA), lysed by sonication in the presence of SERVA protease (50 ug/mL), AEBSF protease (50 ug/mL) and DNAseI (20 ug/mL) and cleared through ultracentrifugation at 40,000 rpm for 45 minutes at 4°C (Beckman Coulter). Cleared lysate was applied to a 5 mL MBPTrap™ HP (Cytiva) pre-equilibrated in lysis buffer. A 2 CV ATP wash (lysis buffer supplemented with 1 mM ATP) and high salt wash (50 mM HEPES, 800 mM NaCl, 10% glycerol, 1 mM MgCl_2_) was performed before extensive washing in lysis buffer (15 CV). The bound protein was eluted with a lysis buffer containing 10 mM maltose. Due to the instability of FL Red1 protein and low yield following expression, subsequent purification steps were not performed. Instead, fractions containing pure Red1 protein were exchanged into SEC buffer (50 mM HEPES, 300 mM NaCl, 10% glycerol, 1 mM TCEP), concentrated using a 50 kDa MWCO Amicon concentrator, flash frozen and stored at -70°C. Red1 NH_2_-terminal constructs (residues 1-362 WT, CM*), closure motifs (residues 340-362 WT, CM*) and coiled-coil proteins (733-819/827 WT, I743R mutants) were produced as 3C HRV cleavable NH_2_-terminal MBP fusion proteins in BL21 STAR *E. coli* cells. Cell cultures were grown at 37°C shaking at 150 rpm until an OD of 0.6 was achieved. Protein expression was induced by the addition of 250 µM IPTG, and expression continued at 18°C for 18 hr. Harvested cells were resuspended in lysis buffer (50 mM HEPES pH 7.5, 300 mM NaCl, 10% glycerol, 1 mM MgCl_2_, 0.1% Triton-X), and for the Red1 N-terminus constructs, 25 mM arginine and glutamic acid were supplemented to the lysis buffer. Cells were lysed by sonication in the presence of SERVA protease (50 µg/mL), AEBSF protease (50 µg/mL) and DNAseI (20 µg/mL) and lysate was cleared through ultracentrifugation at 35,000 rpm for 45 minutes at 4°C (Beckman Coulter). Cleared lysate was applied to a 5 mL MBPTrap™ HP (Cytiva) pre-equilibrated in lysis buffer. A 2 CV ATP wash (lysis buffer supplemented with 1 mM ATP) and high salt wash (50 mM HEPES, 800 mM NaCl, 10% glycerol, 1 mM MgCl_2_) was performed before extensive washing in lysis buffer (15 CV). The bound protein was eluted with a lysis buffer containing 10 mM maltose, concentrated using either 10 or 30 kDa MWCO Amicon concentrators and loaded on Superdex 200 10/300 pre-equilibrated in SEC buffer. Fractions containing purified MBP-Red1 constructs were concentrated, flash frozen and stored at - 70°C.

#### Hop1-Red1 complex

Baculovirus corresponding to Red1^I743R^-MBP and strep-Hop1 were generated through three rounds of amplification in SF9 cells grown in Sf-900 III media (ThermoFisher), shaking 150 RPM at 27°C. For protein expression, Hi5 cells were co-infected with the amplified Hop1/Red1 baculovirus at a ratio of 1:100 (v/v ratio), and cells were cultured for 72 hrs post infection. Harvested cells were washed with PBS, resuspended in lysis buffer (HEPES pH 7.5, 300 mM NaCl, 10% glycerol, 1 mM MgCl_2_, 0.1% Triton-X, 1 mM EDTA), lysed by sonication in the presence of SERVA protease (50 µg/mL), AEBSF protease (50 µg/mL) and DNAseI (20 µg/mL) and cleared through ultracentrifugation at 40,000 rpm for 45 minutes at 4°C (Beckman Coulter). Cleared lysate was applied to a 5 mL Strep-Tactin™XT Superflow™ column (IBA Lifesciences) pre-equilibrated in lysis buffer. A 2 CV ATP wash (lysis buffer supplemented with 1 mM ATP) and high salt (50 mM HEPES, 800 mM NaCl, 10% glycerol, 1 mM MgCl_2_) was performed before extensive washing in lysis buffer (15 CV). The bound protein was eluted with a lysis buffer containing 50 mM biotin and loaded on a HiTrap™ Heparin HP column (GE Healthcare) and subsequently eluted with increasing salt gradient to 1 M NaCl. Fractions containing purified Hop1-Red1 complex were concentrated on a 50 kDa MWCO Amicon concentrator, flash frozen and stored at -70°C.

### Pulldown using purified recombinant protein

#### Hop1^HORMA^ + Red1 ^(1-362)/CM*^ and Red1^CM^ pulldown

Strep-tag pulldown assays were performed with purified strep-Hop1 HORMA domain (bait), MBP-Red1 (1-362) WT/CM* (prey) and MBP-Red1 (340-362) (competitor prey) proteins. 40 µL reactions with 10 µg each Hop1 and Red1 N-terminus constructs in pulldown buffer (20 mM Tris-HCL pH 8.5, 250 mM NaCl, 7.5% glycerol, 1 mM MgCl_2_, 0.05% Tween20) were incubated for 15 minutes at 28°C. Following incubation, 25 µg of MBP-Red1 was titrated in and a further incubation for 10 minutes at 28°C was performed. Samples were added to 5 µL of Strep-Tactin®XT 4Flow® resin preblocked with 1 mg/mL of BSA, and incubated for 15 minutes rotating at 8°C. Beads were washed 3 times with 500 µL of pulldown buffer, before eluting with 30 µL of pulldown buffer supplemented with 10 mM biotin. Input and elution samples were prepared for SDS-PAGE, and gels were visualised with Coomassie staining (Der Blaue Jonas) and Western Blot.

#### Hop1^K593A^ and Red1^CM/CM*^ pulldown

Strep-tag pulldowns assays were performed with purified strep-Hop1 FL K593A (bait) and MBP-Red1 (340-362)/CM*. 40 µL reactions with 10 µg each Hop1 and Red1 in pulldown buffer (50 mM HEPES pH 7.5, 200 mM NaCl, 10% glycerol, 0.05% Tween20) were incubated for 15 minutes at 28°C. Samples were added to 5 uL of Strep-Tactin®XT 4Flow® resin preblocked with 1 mg/mL of BSA, and incubated for 15 minutes rotating at 8°C. Beads were washed 3 times with 500 uL of pulldown buffer, before eluting with 30 µL of pulldown buffer supplemented with 10 mM biotin. Input and elution samples were prepared for SDS-PAGE, and gels were visualised with Coomassie staining (Der Blaue Jonas) and Western Blot.

#### Co-expression pulldowns

For Red1/Mek1 and Red1/Hop1 co-expression pulldowns in SF9 insect cells, Red1 constructs (1-230, 230-345, 230-362, 1-362Δ (230-345), 1-345, 1-362, 1-819, 1-819 I743R, 1-827 and 1-827 I743R) were produced as 3C HRV cleavable N-terminal MBP fusion proteins. Mek1-strep (bait), strep-Hop1 (bait) and MBP-Red1 (prey) baculoviruses were also prepared as stated in the methods section above. For the co-expression, SF9 cells were infected with the amplified bait and prey baculovirus at a ratio of 1:100 (v/v), and cells were cultured for 72 hrs post infection. Harvested cells were washed with PBS, resuspended in lysis buffer (50 mM HEPES pH 7.0, 300 mM NaCl, 7.5% glycerol, 1 mM MgCl2, 0.1% Triton-X), lysed by sonication in the presence of SERVA protease (50 µg/mL), AEBSF protease (50 µg/mL) and DNAseI (20 µg/mL) and cleared through ultracentrifugation at 15,000 rpm for 15 minutes at 4°C (Hettich benchtop centrifuge). The cleared lysate was added to 5 µL of Strep-Tactin®XT 4Flow® resin preblocked with 1 mg/mL of BSA, and incubated for 25 minutes rotating at 8°C. Beads were washed 3 times with 1000 µL of pulldown buffer, before eluting with 30 µL of lysis buffer supplemented with 50 mM biotin. Cleared lysate and elution samples were prepared for SDS-PAGE, and gels were visualised with Coomassie staining (Der Blaue Jonas) and Western Blot.

### Mass Photometry

Mass photometry was performed in 50 mM HEPES pH 7.5, 150 mM NaCl, 5% glycerol, 1 mM TCEP using the RefeynOne mass photometer (Refeyn Ltd., Oxford UK). Thawed proteins were diluted to 30 nM (Hop1-Red1 complex) or 100 nM (Red1 coiled-coil proteins) with the aforementioned buffer immediately before analysis on a glass slide and 1 minute movies were obtained. Peaks were assigned by Gaussian fitting and molecular masses were determined in the Refeyn DiscoverMP software using a NativeMark (Invitrogen) based standard curve as a calibrant under the identical buffer composition.

### SEC-MALS

50 µL samples at 10 µM concentration were loaded onto a Superose6 5/150 analytical size exclusion column (GE Healthcare) equilibrated in buffer containing 10 mM Tris-HCL pH 7.5, 150 mM NaCl, 20 µM ZnSO_4_, 1 mM TCEP attached to an 1260 Infinity II LC System (Agilent). MALS was carried out using a Wyatt DAWN detector attached in line with the size exclusion column. For the analysis, the baseline was manually adjusted and peaks were selected using the built in software (Astra7).

### Cross-linking mass spectrometry

For cross-linking mass spectrometry, proteins were dissolved in 200 ul of buffer (30 mM HEPES pH 7.5, 1 mM TCEP, 300 mM NaCl) to final concentration of 3 μM, mixed with 3 μl of 200 mM DSBU and incubated at 25°C for 1 hr. The reaction was quenched by addition of 20 μl of 1 M Tris pH 8.0 and incubated at 25°C for 30 min. The crosslinked sample was precipitated by the addition of 4X volumes of 100% cold acetone overnight in -20°C and subsequently analyzed as previously described ^86^.

## Acknowledgements

We thank Andrea Musacchio (Max Planck Institute of Molecular Physiology, Dortmund, Germany) for support, Christopher Rath (Max Planck Institute of Molecular Physiology, Dortmund, Germany) for technical support, Franziska Müller and Petra Janning (Max Planck Institute of Molecular Physiology, Dortmund, Germany) for the generation of XL-MS data, Susanna Astrinidis, Rahmiye Kürkcü and Jennifer Jüngling for cloning assistance in the JRW lab. Sharvari Tendulkar and Stefan Westermann (University of Duisburg-Essen, Duisburg, Germany) for experimental advice and help, and Michael Chang (ERIBA, Groningen, The Netherlands) for sharing yeast strains. Members of the Vader lab are acknowledged for insightful discussions and ideas, with special thanks to Richard Cardoso da Silva. We thank Maud Schoot Uiterkamp (Princess Máxima Center for pediatric oncology, Utrecht, The Netherlands) for comments on the manuscript. This work was made possible by financial support from the Max Planck Society, the European Research Council (ERC-StG URDNA, agreement nr. 638197, to GV), the Amsterdam UMC (Amsterdam UMC Research Fellowship 2020, to GV), and the German Research Foundation (WE 6513/2-1 to JRW).

## Author contributions

Conceptualization: GV and JRW; Experimentation: LC, VNN. and SF; Formal analysis: GV, JRW, LC, VNN, and SF; Data curation: GV, JRW, LC, VNN, and SF; Writing—original draft: GV and JRW; Writing—review and editing: GV, JRW, LC, and VNN; Visualization: GV and JRW; Supervision: GV and JRW; Project administration: GV and JRW; Funding acquisition: GV and JRW.

## Correspondence & Materials

Correspondence and requests for materials: GV and JRW.

