## Supplemental Figures 1-7, Supplemental Tables 1-2 for "Organizational principles governing assembly and activation of the meiosis-specific Red1-Hop1-Mek1 complex"

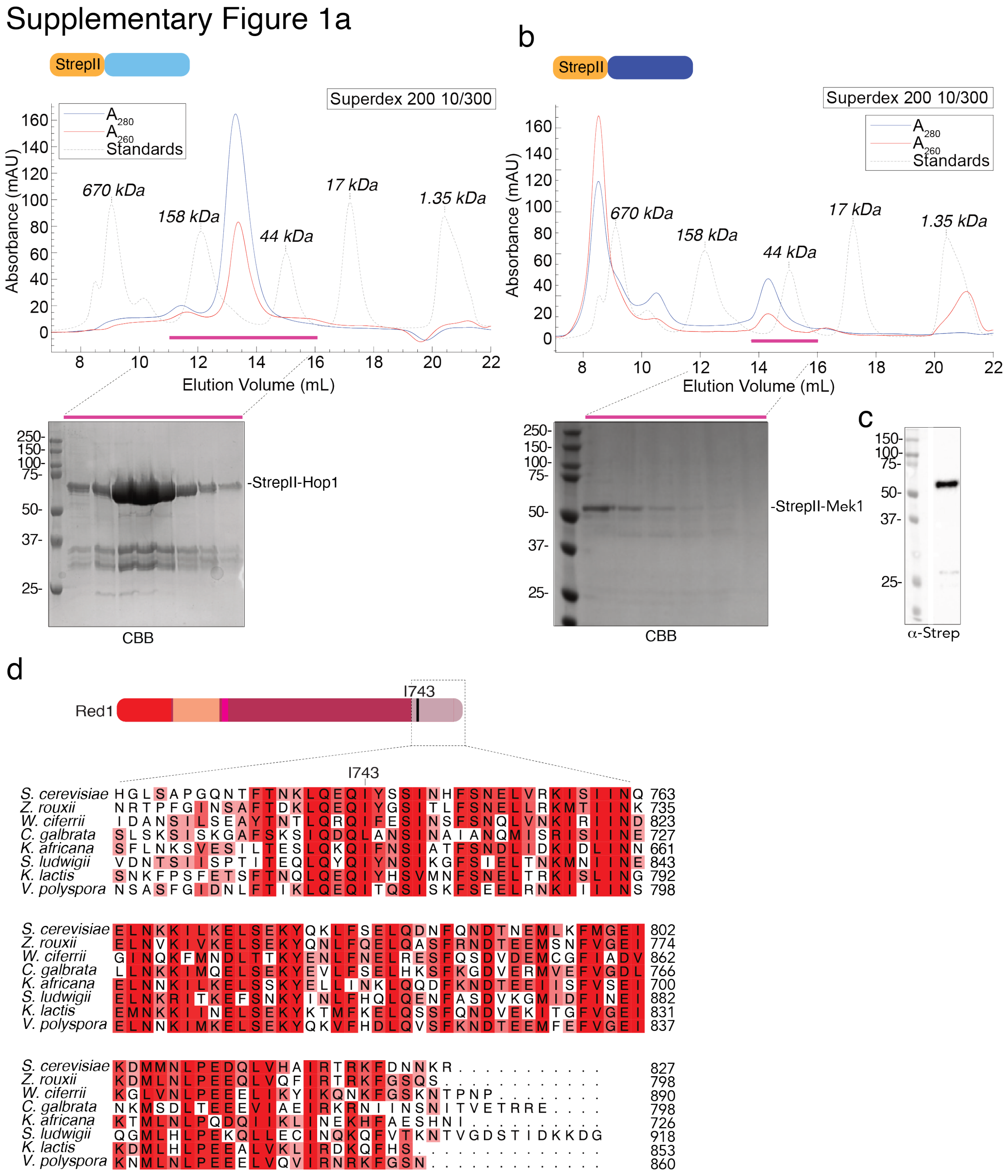


**Supplementary Figure 1.**

**a**. SEC run, including SDS-PAGE, of StrepII-Hop1. CBB = coomassie brilliant blue. b. SEC run, including SDS-PAGE, of StrepII-Mek1. CBB = coomassie brilliant blue. c. SDS-PAGE/western blot of purified StrepII-Mek1 with α-Strep. d. Previously identified Red1 orthologs from different budding yeasts [78] were aligned using MUSCLE, focusing on the COOH-terminal region including the reported tetramerization/filament forming regions [13]. The position of the I743R mutation is shown.

Supplementary Figure 2a

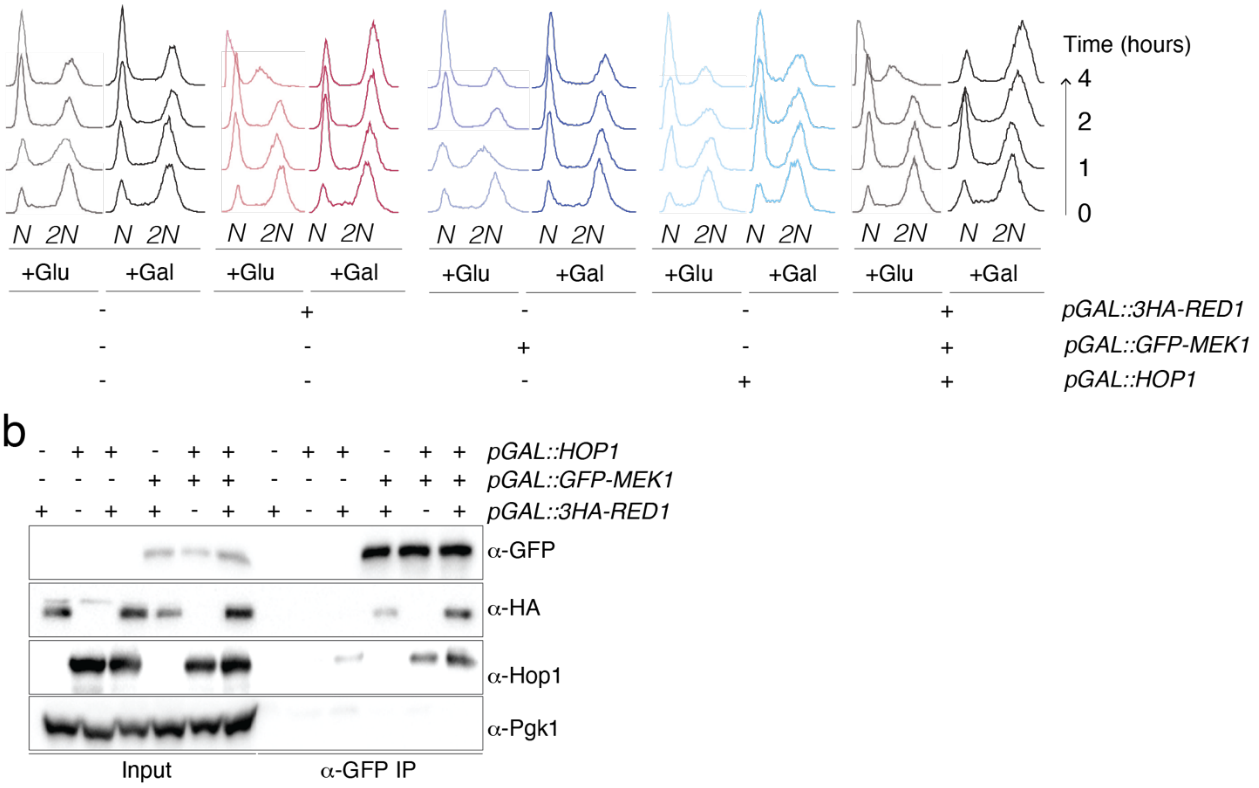


**Supplementary Figure 2.**

**a**. Flow cytometry of wild type, *pGAL::3HA-RED1*, *pGAL::HOP1*, *pGAL::GFP-MEK1* and *RED1*, *HOP1* and *MEK1* strains, upon growth in glucose (+Glu) or galactose-containing (+Gal) conditions. Times (hours) indicated. DNA was stained with SYTOX Green. Strains used are: yGV104, yGV3726, yGV3243, yGV2812, and yGV4806. **b**. Co-immunoprecipitation (co-IP) between Mek1, Hop1 and/or Red1. Mek1 was immunoprecipitated via α-GFP pulldown. α-HA was used to detect Red1, α-GFP was used to detect Mek1, and α-Hop1 was used to detect Hop1. Pgk1 was probed as loading control. Samples were taken after 4 hours induction with galactose. The following strains were used: yGV3242, yGV3726, yGV3235, yGV3255, yGV3219, yGV4806

#### Supplementary Figure 3a

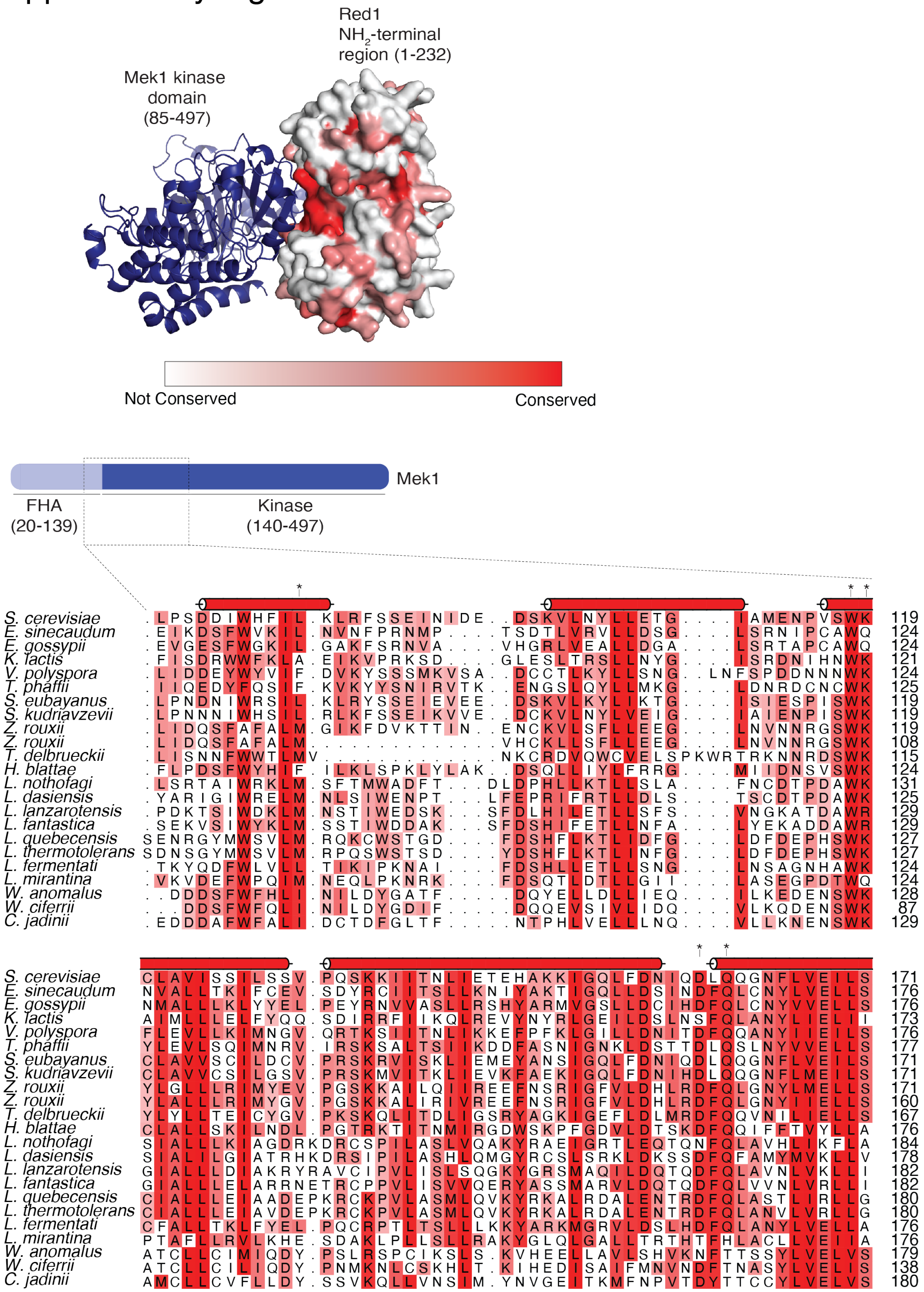


 **Supplementary Figure 3.**

**a.** Previously identified Red1 orthologs from different budding yeasts [78] were aligned using MUSCLE. A subset of these sequences were then used to generate a MSA that was mapped onto the surface of Red1 1-232 using ConSurf [119]. Sequences were coloured in shades of red according to conservation as shown in B). Note the deep red patch that constitutes a hypothetical Mek1 binding site. **b.** Selected region of Red1 MSA described in **a**. and secondary structure for *S. cerevisiae* sequence based upon the AlphaFold2 model. The putative interaction residues for Mek1 are highlighted with *.

Supplementary Figure 4a

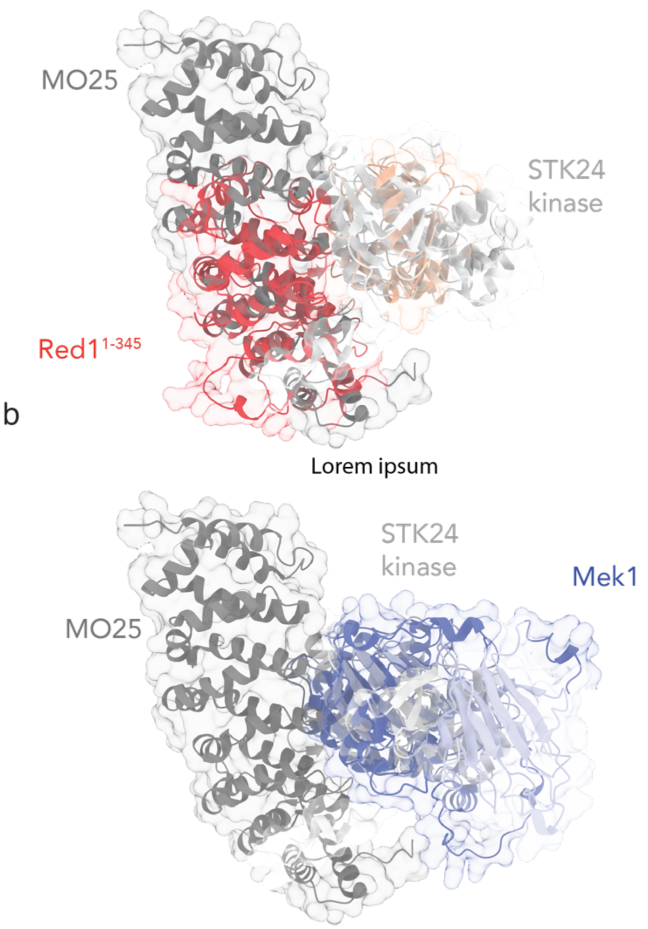


**Supplementary Figure 4.**

**a**. Superposition of the MO25-STK24 crystal structure (3ZHP) on the Red1$^{ARML}$ domain that was discovered via a DALI search (Z-score 9.6, RMSD 3.0Å over 155 residues). The MO25 is shown in dark grey, the STK24 kinase in light grey. b. Matchmaker superposition of the AF2 model of Mek1 onto STK24 kinase (sequence alignment score = 417.8 RMSD between 130 pruned atom pairs is 1.035 angstroms; (across all 263 pairs: 5.928)).

### Supplementary Figure 5a

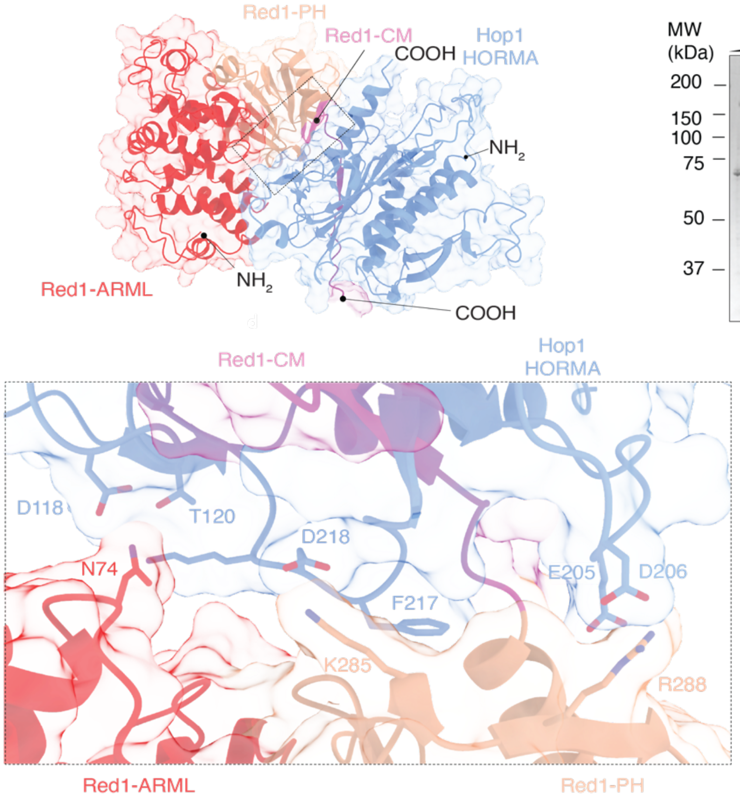


d

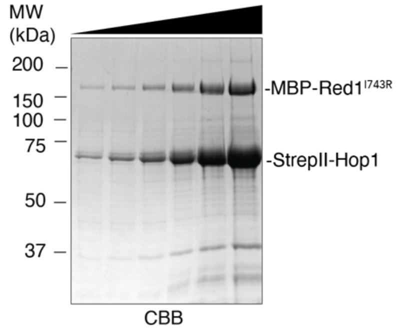


b

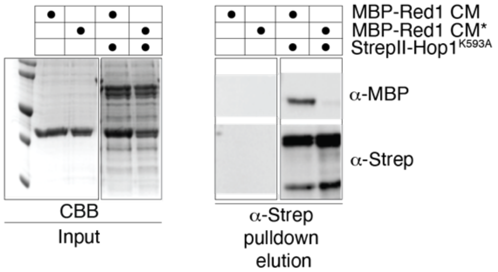


c

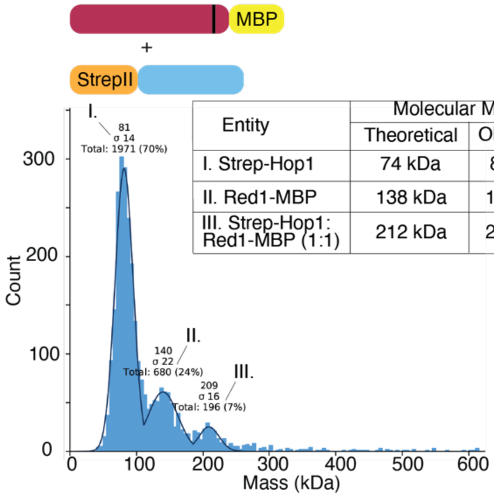


| Entity | Molecular Mass | |
|---|---|---|
| | Theoretical | Observed |
| I. Strep-Hop1 | 74 kDa | 81 kDa |
| II. Red1-MBP | 138 kDa | 140 kDa |
| III. Strep-Hop1: Red1-MBP (1:1) | 212 kDa | 209 kDa |

**Supplementary Figure 5.**

**a**. AlphaFold-based modeling of Red1$^{\text{ARML-PH}}$-Hop1$^{\text{HORMA}}$ association, including zoom-in of interaction interfaces between HOP1$^{\text{HORMA}}$ and Red1$^{\text{ARML-PH}}$. **b**. Pulldown experiments (α-Strep-based) of Hop1 together with indicated Red1 truncations/mutants. α-Strep was used to detect Hop11, and α-MBP was used to detect Red1 fragments. CBB = coomassie brilliant blue. **c.** Mass Photometry of Hop1-Red1 complex. The purified Hop1-Red1 complex was diluted to ~30 nM and measured using a Refyen One mass photometer as per the manufacturer's instructions. **d.** Purity of Hop1-Red1 complex. Increasing quantities of the StrepII-Hop1/Red1$^{\text{1743R}}$-MBP complex were run on a 10% SDS-PAGE gel. The gel was subsequently stained with CBB (coomassie brilliant blue). and imaged.

Supplementary Figure 6a

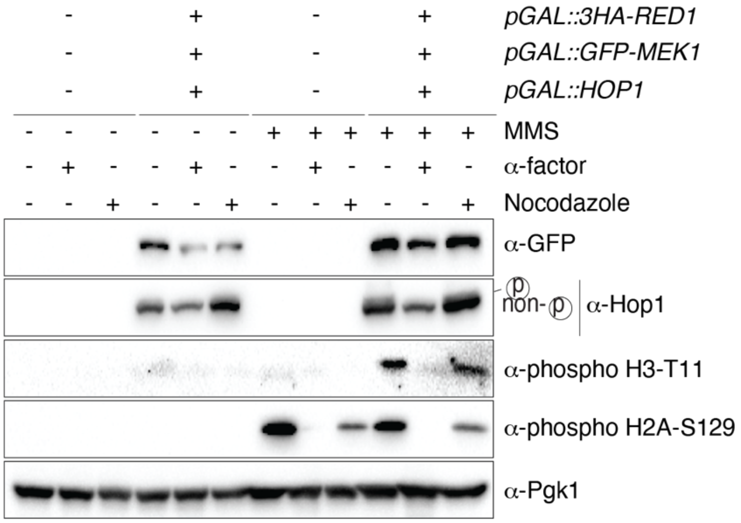


**Supplementary Figure 6.**

**a.** Analysis of Mek1 activation in wild type or Red1, Hop1 and Mek1 expressing cells under different cell cycle conditions. Yeast strains used are yGV104 and yGV480. Cells were arrested in G1-phase by the addition of α-factor, and in mitosis by addition of nocodazole (see Material and Methods for details). MMS was used to induce DNA damage. Galactose-based induction was for 4 hours. α-phospho-H2A-S129 was used to detect Mec1/Tel1-dependent activation, α-phospho-Histone H3-T11 was used to detect Mek1 activation. α-GFP was used to detect Mek1, α-Hop1 was used to detect Hop1 (note also the slower migrating band of Hop1, indicating phosphorylation-mediated gel retardation). Pgk1 was probed as loading control.

#### Supplementary Figure 7a

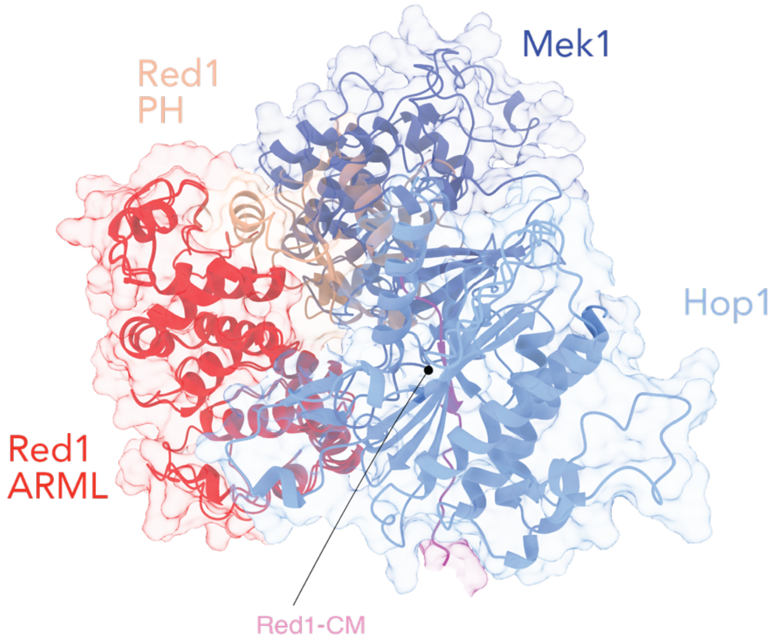


**Supplementary Figure 7.**

**a**. AlphaFold-based modeling of Red1$^{ARML-PH}$-Hop1$^{HORMA}$-Mek1 association.

**Supplemental Table 1. Yeast strains.**

All strains are of W303 background, except yGV49 and yGV4442 which are of the SK1 background.

| **Strain number** | **Genotype** |
|---|---|
| yGV49 | *MATa/MATalpha, ho::LYS2, lys2, ura3, leu2::hisG, his4B::LEU2, ARG4/ arg4-Bgl II* |
| yGV104 | *MATa, ade2-1, leu2-3, ura3, trp1-1, his3-11,15, can1-100, GAL, psi+,* |
| yGV2774 | *MATa, ade2-1, leu2-3, ura3, trp1-1, his3-11,15, can1-100, GAL, psi+, mek1:TRP1-PGAL1-GST-MEK1* |
| yGV2812 | *MATa, ade2-1, leu2-3, ura3, trp1-1, his3-11,15, can1-100, GAL, psi+, mek1:TRP1-PGAL1-GFP-MEK1* |
| yGV3219 | *MATa, ade2-1, leu2-3, ura3, trp1-1, his3-11,15, can1-100, GAL, psi+, Hop1::His3MX6-PGAL1-HOP1, mek1:TRP1-PGAL1-GFP-MEK1* |
| yGV3235 | *MATa, ade2-1, leu2-3, ura3, trp1-1, his3-11,15, can1-100, GAL, psi+, red1::KANMX6::pGAL1-3HA::RED1, hop1::His3MX6-PGAL-HOP1* |
| yGV3243 | *MATa, ade2-1, leu2-3, ura3, trp1-1, his3-11,15, can1-100, GAL, psi+, hop1::His3MX6-PGAL1-HOP1* |
| yGV3255 | *MATa, ADE2, leu2-3, ura3, trp1-1, his3-11,15, can1-100, GAL, psi+, red1::KANMX6::pGAL1-3HA::RED1, mek1:TRP1-PGAL1-GFP-MEK1* |

| | |
|---|---|
| yGV3605 | *MATa, ade2-1, leu2-3, ura3, trp1-1, his3-11,15, can1-100, GAL, psi+, sml1Δ::HphMX6* |
| yGV3693 | *MATa, ade2-1, leu2-3,112, ura3-1, trp1-1, his3-11,15, can1-100,RAD5, mec1Δ::TRP1, sml1Δ::HIS3* |
| yGV3719 | *MATa, ade2-1, leu2-3,112, ura3-1, trp1-1, his3-11,15, can1-100, tel1Δ::URA3* |
| yGV3726 | *MATa, ade2-1, leu2-3,112, ura3-1, trp1-1, his3-11,15, can1-100, red1::KANMX6::pGAL-3HA::RED1* |
| yGV3753 | *MATa, ade2-1, leu2-3,112, ura3-1, trp1-1, his3-11,15, can1-100, rad51Δ::NATMX* |
| yGV3798 | *MATa, ade2-1, leu2-3, ura3, trp1-1, his3-11,15, can1-100, GAL, psi+, red1::KANMX6::pGAL1-3HA-345-827-red1* |
| yGV3799 | *MATa, ade2-1, leu2-3, ura3, trp1-1, his3-11,15, can1-100, GAL, psi+, red1::KANMX6::pGAL1-3HA-367-827-red1* |
| yGV4190 | *MATa, ade2-1, leu2-3, ura3, trp1-1, his3-11,15, can1-100, GAL, psi+, red1::KANMX6::pGAL1-3HA-1-730-red1::TRP1* |
| yGV4191 | *MATa, ade2-1, leu2-3, ura3, trp1-1, his3-11,15, can1-100, GAL, psi+, red1::KANMX6::pGAL1-3HA-1-818-red1::TRP1* |

| | |
|---|---|
| yGV4193 | *MATa, ade2-1, leu2-3, ura3, trp1-1, his3-11,15, can1-100, GAL, psi+, red1::KANMX6::pGAL1-3HA-1-346-red1::TRP1* |
| yGV4194 | *MATa, ade2-1, leu2-3, ura3, trp1-1, his3-11,15, can1-100, GAL, psi+, red1::KANMX6::pGAL1-3HA-1-367-red1::TRP1* |
| yGV4207 | *MATalpha, ade2-1, leu2-3, ura3, trp1-1, his3-11,15, can1-100, GAL, psi+, red1::KANMX6::pGAL1-3HA-345-827-red1, hop1::His3MX6-PGAL1-Hop1, mek1:TRP1-PGAL1-GFP-MEK1* |
| yGV4393 | *MATalpha, ade2-1, leu2-3, ura3, trp1-1, his3-11,15, can1-100, GAL, psi+, hop1::His3MX6-PGAL1-Hop1, mek1:TRP1-PGAL1-GFP-MEK1, red1::KANMX6::pGAL1-3HA-1-818-red1::TRP1* |
| yGV4395 | *MATalpha, ade2-1, leu2-3, ura3, trp1-1, his3-11,15, can1-100, GAL, psi+, Hop1::His3MX6-PGAL1-Hop1, mek1:TRP1-PGAL1-GFP-MEK1, red1::KANMX6::pGAL1-3HA-1-367-red1::TRP1* |
| yGV4397 | *MATa, ade2-1, leu2-3, ura3, trp1-1, his3-11,15, can1-100, GAL, psi+, Hop1::His3MX6-PGAL1-Hop1, mek1:TRP1-PGAL1-GFP-MEK1, red1::KANMX6::pGAL1-3HA-1-346-red1::TRP1* |
| yGV4400 | *MATalpha, ade2-1, leu2-3, ura3, trp1-1, his3-11,15, can1-100, GAL, psi+, Hop1::His3MX6-PGAL1-Hop1, mek1:TRP1-PGAL1-GFP-MEK1, red1::KANMX6::pGAL1-3HA-1-730-red1::TRP1* |

| | |
|---|---|
| yGV4402 | *MATa, ade2-1, leu2-3, ura3, trp1-1, his3-11,15, can1-100, GAL, psi+, hop1::His3MX6-PGAL1-Hop1, mek1:TRP1-PGAL1-GFP-MEK1, red1::KANMX6::pGAL1-3HA-367-827-red1::TRP1* |
| yGV4442 | *MATa/ MATalpha, ho::LYS2, lys2, ura3, leu2::hisG, TRP1, his4B::LEU2, MEK1-6HA::KanMX6, hop1::LEU2* |
| yGV4806 | *MATa, ade2-1, leu2-3, ura3, trp1-1, his3-11,15, can1-100, GAL, psi+, red1::KANMX6::pGAL1-3HA::RED1, Hop1::His3MX6-PGAL1-Hop1, mek1:TRP1-PGAL1-GFP-MEK1* |
| yGV5011 | *MATa, ade2-1, leu2-3,112, ura3-1, trp1-1, his3-11,15, can1-100, GAL, psi+,red1::KANMX6::pGAL1-3HA::RED1, Hop1::His3MX6-PGAL1-HOP1, mek1:TRP1-PGAL1-GFP-MEK1, tel1Δ ::URA3* |
| yGV5033 | *MATa, ade2-1, leu2-3,112, ura3-1, trp1-1, his3-11,15, can1-100,RAD5, mek1:TRP1-PGAL1-GFP-MEK1, red1::KANMX6::pGAL1-3HA::RED1, Hop1::His3MX6-PGAL1-HOP1, mec1Δ ::TRP1, sml1Δ::HIS3* |
| yGV5044 | *MATa, ade2-1, leu2-3, ura3, trp1-1, his3-11,15, can1-100, GAL, psi+, sml1Δ::HphMX6,red1::KANMX6::pGAL-3HA::RED1, Hop1::His3MX6-PGAL-HOP1, mek1:TRP1-PGAL1-GFP-MEK1* |

**Supplemental Table 2. Plasmids.**

| Plasmid ID | Description | Insert | Expression system | Affinity tag(s) | Reference |
|---|---|---|---|---|---|
| pWL658 | StrepII-Hop1 | Hop1 full length (1-605) | Bacteria | $NH_2$-terminal StrepII | [15] |
| pWL661 | StrepII-Hop1 | Hop1 full length (1-605) | Insect cells | $NH_2$-terminal StrepII | [15] |
| pWL1139 | StrepII-Hop1$^{K593A}$ | Hop1 full length /K593A | Bacteria | $NH_2$-terminal StrepII | [15] |
| pWL1375 | StrepII-Hop1$^{1-255}$ | Hop1 HORMA domain (1-255) | Bacteria | $NH_2$-terminal StrepII | [15] |
| pWL432 | MBP-Red1 | Red1 full length (1-827) | Insect cells | $NH_2$-terminal MBP | This study |
| pWL543 | MBP-Red1$^{I743R}$ | Red1 full length (1-827)/I743R | Insect cells | $NH_2$-terminal MBP | This study |
| pWL1157 | Red1$^{I743R}$-MBP | Red1 full length (1-827)/I743R | Insect cells | COOH-terminal MBP | This study |
| pWL2329 | MBP-Red1$^{1-362}$ | Red1 1-362 (AMRL-PH-CM) | Bacteria | $NH_2$-terminal MBP | This study |
| pWL482 | MBP-Red1 $^{1-362}$ | Red1 1-362 (AMRL-PH-CM) | Insect cells | $NH_2$-terminal MBP | This study |
| pWL2458 | MBP-Red1$^{1-362/CM*}$ | Red1 1-362 (AMRL-PH)/ CM* | Bacteria | $NH_2$-terminal MBP | This study |
| pWL484 | MBP-Red1$^{1-345}$ | Red1 1-345 (AMRL-PH) | Insect cells | $NH_2$-terminal MBP | This study |
| pWL2247 | MBP-Red1$^{1-230}$ | Red1 1-230 (AMRL) | Insect cells | $NH_2$-terminal MBP | This study |
| pWL2249 | MBP-Red1$^{230-344}$ | Red1 230-344 (PH) | Insect cells | $NH_2$-terminal MBP | This study |
| pWL2154 | MBP Red1$^{230-362}$ | Red1 230-362 (PH-CM) | Insect cells | $NH_2$-terminal MBP | This study |
| pWL372 | MBP-Red1$^{340-362}$ | Red1 340-362 (CM) | Bacteria | $NH_2$-terminal MBP | This study |

| | | | | | |
|---|---|---|---|---|---|
| pWL2459 | MBP-Red1$^{340-362/CM*}$ | Red1 340-362 (CM)/CM* | Bacteria | $NH_2$-terminal MBP | This study |
| pWL542 | MBP-Red1$^{1-819}$ | Red1 1-819 | Insect cells | $NH_2$-terminal MBP | This study |
| pWL2384 | MBP-Red1$^{1-819/I743R}$ | Red1 1-819/I743R | Insect cells | $NH_2$-terminal MBP | This study |
| pWL960 | MBP-Red1$^{733-827}$ | Red1 733-827 | Bacteria | $NH_2$-terminal MBP | This study |
| pWL2369 | MBP-Red1$^{733-827/I743R}$ | Red1 733-827/I743R | Bacteria | $NH_2$-terminal MBP | This study |
| pWL2363 | MBP-Red1$^{733-819}$ | Red1 733-819 | Bacteria | $NH_2$-terminal MBP | This study |
| pWL2364 | MBP-Red1$^{733-819/I743R}$ | Red1 733-819/I743R | Bacteria | $NH_2$-terminal MBP | This study |
| pWL425 | Mek1-StrepII | Mek1 full length (1-497) | Insect cells | COOH-terminal StrepII | This study |
